# Selection for non-specific adhesion is a driver of FimH evolution increasing *Escherichia coli* biofilm capacity

**DOI:** 10.1101/2021.08.03.454899

**Authors:** Mari Yoshida, Stanislas Thiriet-Rupert, Leonie Mayer, Christophe Beloin, Jean-Marc Ghigo

## Abstract

Bacterial interactions with surfaces rely on the coordinated expression and interplay of surface exposed adhesion factors. However, how bacteria dynamically modulate their vast repertoire of adhesins to achieve surface colonization is not yet well-understood. We used experimental evolution and positive selection for improved adhesion to investigate how an initially poorly adherent *Escherichia coli* strain increased its adhesion capacities to abiotic surfaces. We showed that all identified evolved clones acquired mutations located almost exclusively in the lectin domain of *fimH*, the gene coding for the *α*-D-mannose-specific tip adhesin of type 1 fimbriae. While most of these *fimH* mutants showed reduced mannose-binding ability, they all displayed enhanced binding to abiotic surfaces, indicating a trade-off between FimH-mediated specific and non-specific adhesion properties. Several of the identified mutations were already reported in FimH lectin domain of pathogenic and environmental *E. coli,* suggesting that, beyond patho-adaptation, FimH microevolution favoring non-specific surface adhesion could constitute a selective advantage for natural *E. coli* isolates. Consistently, although *E. coli* deleted for the *fim* operon still evolves an increased adhesion capacity, mutants selected in the Δ*fim* background are outcompeted by *fimH* mutants revealing clonal interference for adhesion. Our study therefore provides insights into the plasticity of *E. coli* adhesion potential and shows that evolution of type 1 fimbriae is a major driver of the adaptation of natural *E. coli* to colonization.

## INTRODUCTION

Adhesion and subsequent formation of biofilms on environmental or host surfaces enable bacteria to withstand natural mechanical fluxes and is an essential step of most colonization or infection processes ^1–5^. In commensal and pathogenic *Escherichia coli*, adhesion and biofilm formation is achieved using a large arsenal of adhesion factors ^2, 4^, including proteinaceous adhesins protruding from the bacterial cell envelope and promoting non-specific surface adhesion (e.g. amyloid curli), afimbrial autotransporter adhesins mediating aggregation via homotypic self-interactions (e.g. Ag43 adhesin) or polymeric fimbrial adhesins that specifically recognize oligosaccharidic ligands (e.g. type 1 fimbriae) ^6–10^. *E. coli* also secretes several polysaccharidic exopolymers such as cellulose, ß-1,6-*N*-acetyl-D-glucosamine polymer and capsular polysaccharides, promoting or hindering adhesion on living or inert surfaces ^11–15^. In addition to these well characterized surface structures, the *E. coli* genome contains several partially characterized genes or operons encoding potential adhesion factors ^6, 7, 16, 17^. The expression of *E. coli* adhesins is controled by diverse regulatory networks enabling rapid transcriptional change in response to stresses and environmental cues such as oxygen levels, pH and chemical gradients ^6, 18–20^. However, how *E. coli* controls its adhesive properties and coordinates the interplay between its adhesion factors, while avoiding antagonistic physical interferences between adhesins is still poorly understood.

Here, we used a dynamic *in vitro* biofilm model amenable to experimental evolution to explore how *E. coli* adapts when subjected to positive selection for increased capacity to bind to an abiotic surface and form biofilms. We showed that, despite the diversity of *E. coli* adhesins, all end-point populations displaying increased biofilm biomass after positive selection almost exclusively acquired mutations in *fimH*, the gene coding for the *α*-D- mannose-specific tip adhesin of type 1 fimbriae enabling adhesion to mannosylated epithelial cells. The identified *fimH* mutations, including mutations found in environmental and clinical *E. coli* isolates, displayed enhanced capacity for initial adhesion to abiotic surfaces but reduced mannose-binding properties and outcompeted the strongest biofilm-forming mutant emerging from a selection performed in a strain lacking the whole *fim* operon. This indicates that selection of *fimH* mutants with increased non-specific adhesion capacity could provide commensal or pathogenic *E. coli* a selective advantage for surface colonization and persistence in the environment. Our study therefore provides direct insights into the plasticity of the *E. coli* adhesion repertoire and shows that biofilm formation is a powerful driver of the evolution of *E. coli* adhesion potential.

## RESULTS

### Positive selection of *Escherichia coli* mutants with increased biofilm capacity

We subjected the poor biofilm-forming strain *E. coli* MG1655 to positive selection for spontaneous mutants with increased adhesion capacity using continuous-flow biofilm microfermenters (Fig. 1A). We conducted 12 parallel evolution experiments with 15 cycles of 8 hours biofilm formation followed by overnight planktonic growth (15 days). The capacity of biofilm formation in microfermenters was enhanced for 6 of the 12 evolved population, with a 10 to 100-fold increase for 4 of them as compared to the WT ancestor (G6, R4, R5, R6) (Fig. 1BC and Supp. Fig. S1A). These 6 populations also displayed an enhanced capacity to form biofilms in microtiter plate (Supp. Fig S1B) showing that the evolved trait was robust over two different biofilm models. To study the dynamic of the evolution of the adhesion capacities of the different population we performed microtiter plate biofilm assays for each cycle of the different population (Fig. 1D). Biofilm formation started to increase at cycle 5 for the G6 population and around cycle 10 for the population G1, G3, R4, R5 and R6 (Fig. 1D). Consistently, the enhanced biofilm formation evolved trait was also observed at clones level since heterogeneous but significant increase in biofilm capacity compared to the ancestral WT strain was shown in 15 individual clones isolated from each of the 6 more adherent populations (Fig. 1E and Supp. Fig.S1B).

**Figure 1.**
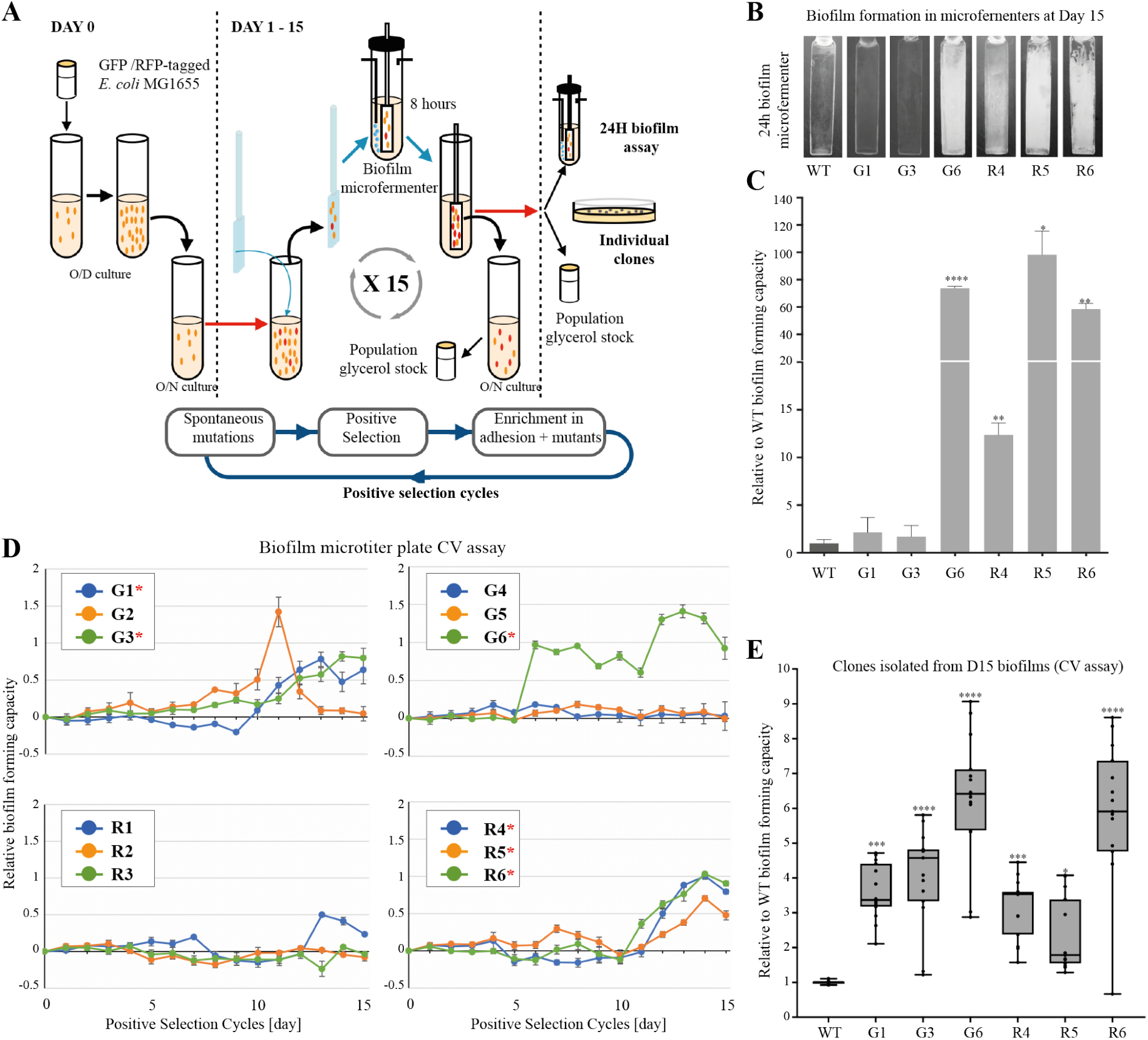
Selection of *E. coli* mutants with increased biofilm capacity. **A.** Schematics of positive selection to identify GFP- or RFP-tagged *E. coli* MG1655 mutants with increased biofilm formation capacity. **B.** Comparison of biofilm formation on biofilm microfermenters spatula of end-point populations. **C.** Comparison of biofilm biomass of end- point populations and controls in biofilm microfermenters. The biofilm biomasses were collected from the spatula incubated for 24 hours and the biofilm-forming capacities were calculated relative to the one obtained with the control ancestral WT (set to 1). **D.** Evolution of biofilm forming capacity of bacterial population subjected to positive selection monitored by performing crystal violet biofilm assay using an aliquot of the population produced at each selection cycle. Comparison with the parental wild type strain and normalization were performed by calculating relative value using WT *E. coli* MG1655 (poor biofilm former) and *E. coli* TG1 strains as follows: (OD_test_-OD_WT_)/(OD _TG1_-OD_WT_). When the value is equal to 1 and 0, the capacity is as same as the TG1 strain and the WT MG1655 parental strain, respectively. The experiments were performed in triplicate. G1-6 populations derive from parental GFP-tagged MG1655. R1-6 populations derive from parental RFP-tagged MG1655. The selected populations with increased biofilm capacity are indicated with an asterisk. **E.** Comparison of biofilm forming capacities of 15 clones isolated from the selected population at cycle 15 by crystal violet biofilm assay. Each point represents the data of an individual clone. Each solid line in the boxplot shows the median of the biofilm forming capacities, and the boxes illustrate the first and the third quartiles. The relative biofilm capacities were calculated using the capacities of the parental wild type strain (median set to 1). All crystal violet biofilm assay experiments were conducted with at least 8 replicates. Continuous flow biofilm experiments in microfermenters were performed 3 times (**C)** and statistics correspond to unpaired t-test with Welch’s corrections. In **E** (12 répétitions) statistics correspond to unpaired, non-parametric Mann-Whitney test comparing all conditions to ancestral WT. * p<0.05; ** p<0.01; *** p<0.001; **** p<0.0001.

### End-point biofilm-positive mutants carry mutations in the gene encoding the type 1 fimbriae tip adhesin FimH

To identify the nature of the mutations leading to increased adhesion, we sequenced and compared the ancestral population to the genome of eight populations at the end of 15 selection cycles, including the six biofilm populations with enhanced biofilm forming capacity (G1, G3, G6, R4, R5, R6) and six populations that showed moderate or no biofilm increase (G2, G4, G5, R1, R2 and R3) (Supp. Fig. S1B). We also sequenced four individual clones isolated from each of the six biofilm positive populations and two of the six biofilm negative populations. The populations and clones with enhanced biofilm capacities all carried mutations at high frequencies located in the gene coding for FimH and no *fimH* mutation was detected in the six populations showing no increase in biofilm capacity after 15 cycles of positive selection (Fig. 2A and Supp. Table S1). FimH is the D-mannose specific adhesin of type 1 fimbriae, a major *E. coli* adhesin enabling epithelial cell colonization and shown to be critical for biofilm formation on abiotic surfaces ^21^. FimH is a 300 residue and 31.473 kDa protein consisting of a pilin domain (amino acid 181-294) and a mannose-binding lectin domain (amino acid 23-177) connected through a 4 amino acid linker peptide chain (Fig. 2B).

**Figure 2.**
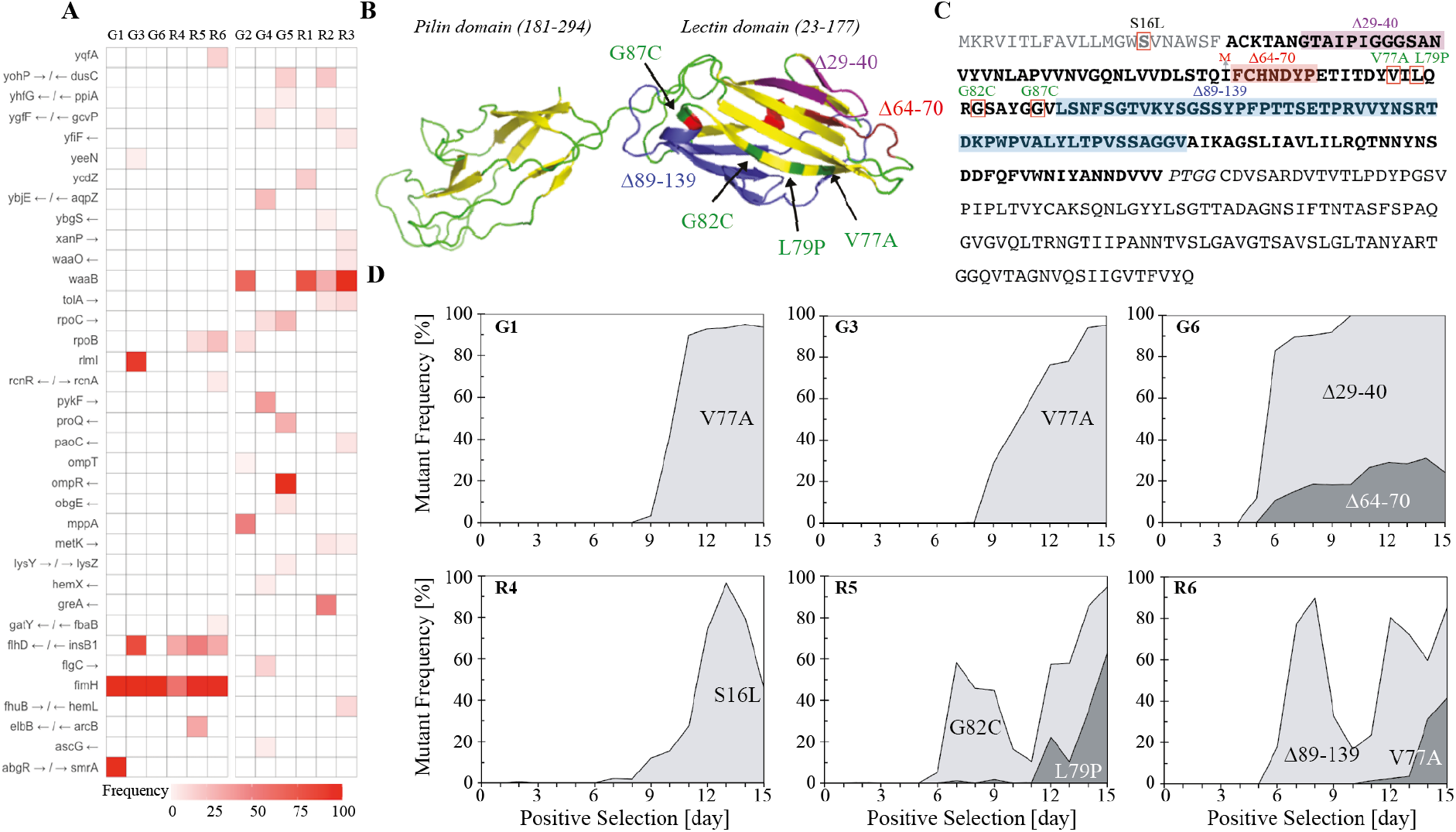
Nature and frequency of FimH mutations identified in evolved populations with increased biofilm capacity. **A.** Frequency of mutation in targeted genes. Frequency of mutations inferred from population sequencing in the 6 populations with increased biofilm formation capacity (left) and the 6 populations without increased biofilm capacity (right). The frequency is color coded in red according to the legend at the bottom of the heatmap and correspond to the total frequency of all mutations in each gene within a population at cycle 15 of the experimental evolution. **B.** Identified mutations in FimH protein structure. The locations of amino acid substitutions are indicated with black arrows and the in-frame deletions are indicated in purple (*Δ*29-40), red (*Δ*64-70) and blue (*Δ*89-139). **C.** Identified FimH mutations are indicated above the FimH amino acid sequence. Single amino acid substitutions are highlighted with red boxes and three deletions are highlighted in pale purple, red and blue. The G87C mutation was identified in a distinct positive selection experiment using less selection cycles (4 instead of 15) and longer residence in the biofilm microfermenters at each cycle (18 hours instead of 8 hours). **D.** Frequency of indicated FimH mutations acquired during the positive selection experiment in populations with increased adhesion capacity (G1, G3, G6, R4, R5, R6). The grey areas show the frequency of the mutants and the white area shows the frequency of the wild type strains. The frequencies were calculated using Sanger sequencing of *fimH* region of sampled populations.

We identified 3 in-frame deletions (Δ29-40 and Δ64-70, and Δ89-139) and three non-synonymous substitutions (V77A, L79P, G82C) in the lectin domain of the FimH protein, as well as one non-synonymous substitution (S16L) in the signal peptide of the FimH protein (Fig. 2BC). Some *fimH* mutations coexisted within populations: L79P and G82C were found in R5 population, V77A and Δ89-139 in R6 population, and Δ29-40 and Δ64-70 in G6 population. By contrast, R4 population only had the S16L substitution, and G1 and G3 populations only had V77A mutations (Supp.Table S1).

In order to check if this outcome was specific to the parameters used in our selection protocol, we ran another experiment using different selection pressures with less selection cycles (4 instead of 15) and longer residence time in biofilm microfermenters at each cycle (18 hours instead of 8 hours). This protocol also resulted in the selection of *fimH* mutation (G87C), emphasizing the predominant role of this adhesin in enhanced binding capacities and biofilm formation (Fig. 2BC). In order to determine the dynamics of emergence and population spread of the identified *fimH* mutations over time, we performed a Sanger sequencing analysis of *fimH* region PCR products of populations collected at each of the 15 positive selection cycles that led to increased biofilm capacities (Fig. 2D). All *fimH* in-frame deletions and the G82C mutation emerged as early as the fourth/fifth cycle, while the other non-synonymous FimH substitutions emerged at cycle 6 or later.

### The phenotypic differences between *fimH* mutants are not due to phase variable expression of the *fim* operon

We isolated individual clones carrying an independent mutation in FimH and measured their biofilm capacity using continuous biofilm microfermenters. While most tested evolved clones showed close to wild-type growth rate, three mutations (V77A, L79P and Δ29-40) slightly impacted growth (Supp. Fig. S2AB) and 1:1 competition between these 3 clones and WT confirmed this fitness default (Supp. Fig. S2C). All clones but the one carrying L79P, demonstrated increased biofilm-forming capacity, *i.e.* 1.8 to 113 times more than WT ancestral strain after 24 hours of incubation in microfermenters (Fig. 3A). The mutants encoding FimH deletions Δ64-70 and Δ89-139 displayed the highest biofilm formation, relative to the ancestral WT (Fig. 3B). This enhanced capacity to form biofilms could be linked to phase variable expression of the *fim* operon that enable rapid on/off change of type 1 fimbriae production by inversion of the *fimA* promoter mediated by FimB and FimE recombinases (see *fim* region in Supp. Fig. S3A) ^22–24^. However, a test of the orientation of the *fim* promoter by PCR did not reveal any significant differences in ON/OFF status between wild type parental and the evolved clones. The mutant clones were mainly OFF in planktonic culture and mainly ON in biofilm on microfermenter spatula, while none of the clones or the evolved populations displayed any mutations known to impact the *fim* switch, thus reflecting an enrichment for *fim* ON status in biofilm conditions (Supp. Fig. S3BC).

**Figure 3.**
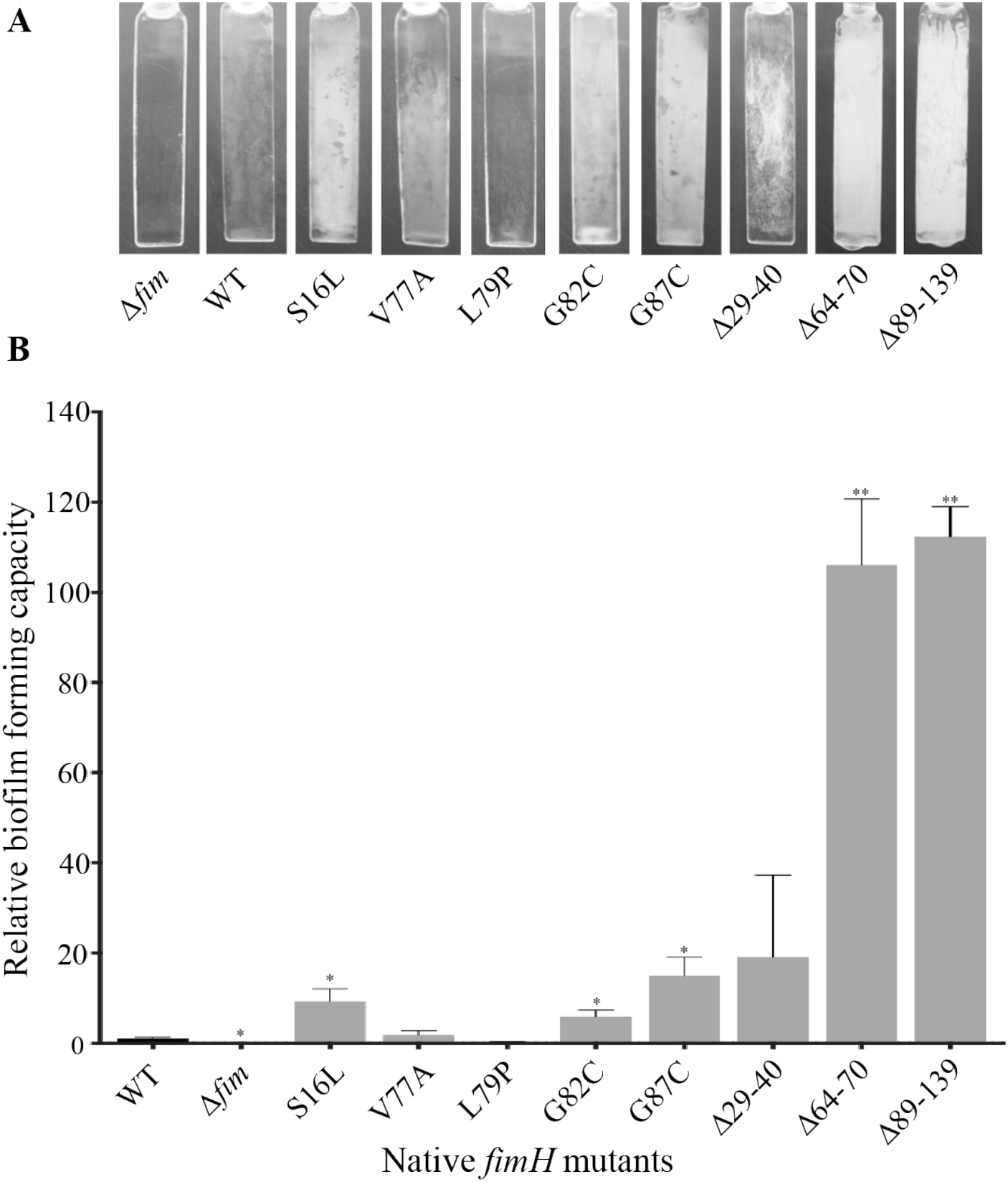
Biofilm formation capacity of individual *fimH* mutants. Biofilm formation of *fimH* mutants was compared after 24 hours incubation in biofilm microfermenters. **A.** Pictures of representative biofilm biomass formed on biofilm microfermenters spatulas**. B.** The relative biofilm-forming capacities of individual *fimH* mutants was determined by comparison with the capacities of the Δ*fim* (0) and wild type strains (1). Microfermenter biofilm experiments were performed in triplicate and statistics correspond to unpaired t-test with Welch’s corrections.. * p<0.05; ** p<0.01.

### Evolved *fimH* mutations increase non-specific initial adhesion

To characterize the phenotypic consequences of the identified *fimH* mutants in planktonic conditions and, considering the ON *fim* status of clones selected in biofilm conditions, we placed the WT and mutant *fim* operon (and therefore corresponding native or mutated *fimH* genes) under the control of the lambda PcL constitutive promoter ^6^. This led to normalized type 1 fimbriae expression, as shown using immunodetection with anti-FimA antibodies (Supp. Fig. S4). Biofilm biomass formed in microfermenters of PcL-*fimH* mutants, compared to PcL-*fimH* WT, showed differences similar to those observed between (non-PcL) mutants and WT strains (compare Fig. 4A with Fig. 3), suggesting that, independently of *fim* phase variation, the selected FimH mutants displayed specific properties enhancing *E. coli* biofilm formation. Interestingly, selected mutants out-competed the wild-type strain for increased initial adhesion to glass in 1:1 competition between WT PcL-*fimH* and mutant PcL-*fimH* (Fig. 4B). This fitness advantage suggests that increased initial non-specific adhesion is one of the main drivers of the enhanced capacity to form biofilms of the *fimH* mutant compared to the wild-type ancestral strain. However, we detected a negative correlation between *fimH* mutants initial adhesion capacity and their biofilm capacity (Spearman’s rho = -0.76, p-value = 0.037, Supp. Fig. S5), suggesting a trade-off between initial adhesion and biofilm maturation.

**Figure 4.**
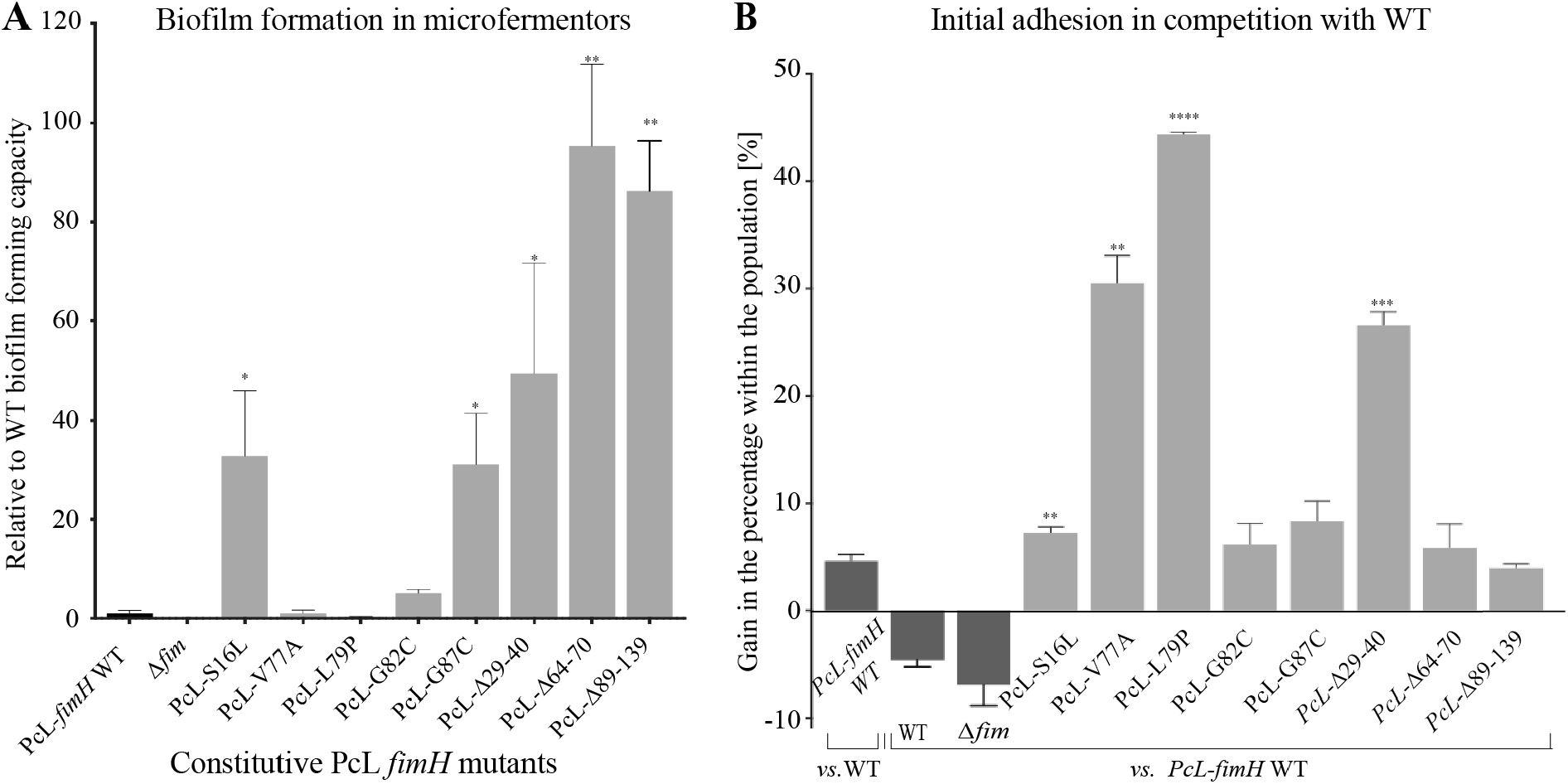
Biofilm and initial adhesion capacity of constitutive PcL *fimH* mutants. **A.** Biofilm formation of *Δfim* and PcL*fimH* WT strains (controls) and individual PcL*fimH* mutants after 24 hours incubation in biofilm microfermenters. The relative biofilm-forming capacities of individual PcL *fimH* mutants was determined by comparison with the capacities of the Δ*fim* (0) and PcL*fimH* WT strains (1). All biofilm experiments were performed in triplicate. **B.** Initial adhesion of PcL*fimH* mutants in competition with PcL*fimH* WT. The PcL*fimH* mutants were inoculated to the microfermenter spatula in 1:1 ratio with PcL*fimH* WT strain carrying either GFP or RFP fluorescent tags for 10 min, followed by washing, resuspension of the attached bacteria and CFU counting. The changes in the percentage of each strain within the population are shown. PcL*fimH* WT is also compared with the other control strains (WT and Δ*fim* strains, and PcL*fimH* WT strain with the different fluorescent tag). Statistics correspond to unpaired t-test with Welch’s corrections. comparing all conditions to ancestral WT. * p<0.05; ** p<0.01; *** p<0.001; **** p<0.0001.

We also evaluated the ability of PcL-*fimH* mutants to bind mannosylated proteins using a yeast agglutination assay performed under static or agitated conditions, where the latter condition promotes catch-bond adhesion ^25^. In both situations, all FimH mutations, except G87C, reduced yeast agglutination capacity, with some mutations displaying almost no agglutination, very similar to a Δ*fimH* strain agglutination phenotype (Supp Fig. S6). This indicates a clear trade-off between FimH-enhanced adhesion to abiotic surfaces and FimH adhesion to mannose.

### Mutations identified in FimH are also found in natural and clinical isolates

In order to assess whether the FimH mutations identified in our *in vitro* evolution experiments reflect evolutionary paths occurring in natural or clinical *E. coli* isolates, we retrieved all *E. coli* FimH sequences available in NCBI protein databank as well as 2067 sequenced genomes of *E. coli* (Supp. Table S2). The resulting NCBI dataset composed of 3266 sequences (Supp. Table S3A) shows a strong bias toward strains of human and animal origin (53.25 % versus 1.81 % for environmental strains, while no information is available for the remaining 45%), likely due to the predominant representation of health-related studies. To partly compensate for this skew, we added the *fimH* sequence of 277 *E. coli* environmental strains from a recent study ^26^ (see Methods section and Supp. Table S3B). When the 3543 total FimH protein sequences extracted from this dataset were compared to MG1655 FimH, 10.4 % (369 FimH sequences) displayed amino-acid changes at the same positions of at least one of the eight mutations identified in our study (Supp. Table S3AB). An enrichment for strains belonging to the B1 phylogroup was identified in these 369 sequences, relative to the whole dataset (Supp. Table S4A). Interestingly, 64 FimH proteins showed mutations that were also identified in our study (one S16L, 18 V77A, one L79P, 42 G87C and two Δ29-40). The G87C mutation (=G66C when amino acid numbering starts after the signal peptide), a position previously reported to be under positive selection in uropathogenic *E. coli* strains, being the most widespread ^27, 28^. These results demonstrate that our study recapitulated some of the selection events driving natural evolution of FimH (Fig. 5A). The enrichment in B1 strains previously identified was not maintained in these 64 sequences, this phylogroup being rather depleted relatively to the whole dataset (Supp. Table S4A). However, there was an enrichment for B2 and F strains (to which most ExPEC strains belong), suggesting that our experimental evolution reflects part of the selective pressures applied to ExPEC strains.

**Figure 5.**
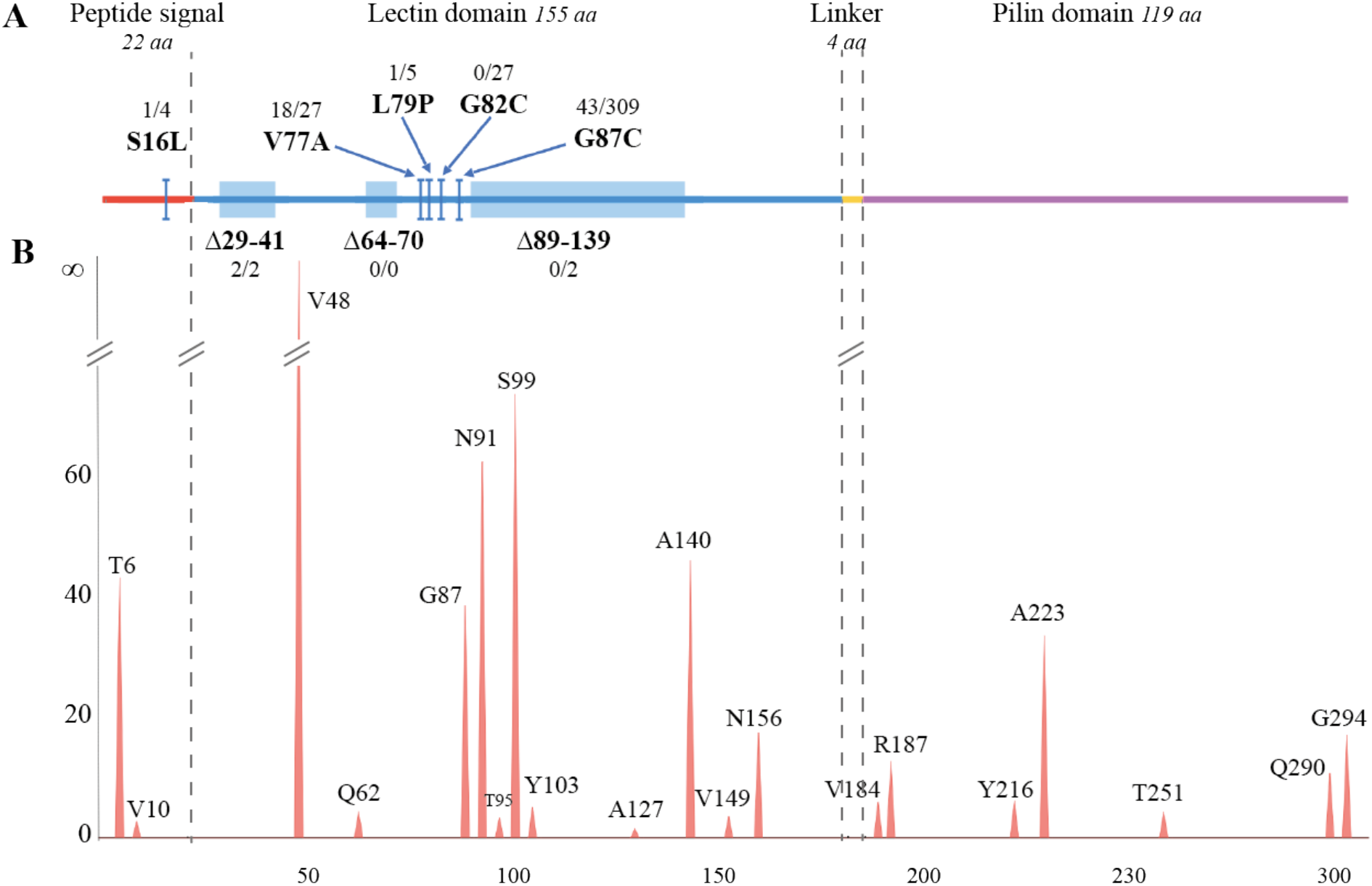
Analysis of the mutational spectrum of FimH sequences. **A.** Linear structure and amino-acid size of FimH protein domains with the eight mutations selected during the experimental evolution. The two numbers provided for each mutation correspond to respectively, the number of mutations among the 3266 analyzed FimH sequences and the number of sequences having a mutation at the same position in this study (regardless of mutation type). **B.** Negative log-transformed significance (-log10(pvalue)) of the mutation frequency at each position of the sequence relatively to the frequency calculated over all sequences from Database sequences. A one-way Anova was performed, the resulting p-values were adjusted and a False Discovery Rate (FDR) of 0.05 was considered as significant. Only positions with a significantly higher mutation frequency are plotted.

### FimH mutational landscape diversity highlights its potential functional plasticity

All selected mutations in our study were found in the FimH lectin domain and none in its pilin domain. To test whether this was a general trend in FimH sequences or rather specific to our experimental settings, we screened all available FimH sequences for single amino-acid changes and calculated their frequency for each structural part of the FimH sequence (the signal peptide, the lectin domain, the four amino-acids linker and the pilin domain). The majority of these mutations were found in the lectin domain and were significantly higher (1.47-fold, One-way Anova: df = 1, p value < 2.2e-16) than expected, while the pilin domain accumulated significantly less mutations (-2.31-fold, One-way Anova: df = 1, p value < 2.2e-16) (Fig. 5B). To analyze the *E. coli* FimH mutational landscape in more details, the same analysis was carried out for each position of the sequence and 20 of them were found to be statistically more prone to mutation (Fig. 5B and Supp. Table S4B). Interestingly, some of the mutations identified at these hotspots were found to co-occur in the same FimH sequences (Supp. Fig. S7). Different phylogroup enrichments were also identified at these polymorphic sites (Supp. Table S4C).

To bring further insight into the evolution of FimH sequences in *E. coli*, we built a phylogenetic tree (Supp. Fig. S8) and analyzed it with the *codeml* program in PAML v4.9 ^29^. For this purpose, we divided the tree into five sub-groups (Sg1 to Sg5) based on phylogroups distribution along the tree (see methods). Overall, the evolutionary analysis showed a prevalence for purifying selection (Supp. Table S5A). However, sequences belonging to Sg3 showed signs of episodic positive selection. Interestingly, most of the sites identified to be positively selected in this group were located in the lectin domain including positions 82 and 87 corresponding to evolved sites in our experimental evolution (Supp. Table S5 AB). Moreover, Sg3 was enriched in strains belonging to phylogroup B2 (Supp. Table S4D) as well as comprised all sequences having amino acids Ser-70 and Asn-78 (Ser-91 and Asn-99 in this study) described to be related to UPEC strains ^27^. Finally, a site partition analysis revealed that, although being both under purifying selection, the lectin and the pilin domains underwent a different evolutionary path in Sg3 and Sg5 (Supp. Table S5CD). Overall, these results demonstrate that the lectin domain of FimH is the most subjected to mutations both in laboratory evolved strains and in natural and clinical isolates, which could reflect complementary selection pressure in environments where bacteria need to withstand natural fluxes and perturbations.

### In absence of *fimH*, evolution towards biofilm formation involves a broader range of mutations

Despite the diversity of *E. coli* K-12 surface structures known to contribute to biofilm formation ^30^, mutations leading to increased biofilm formation revealed by our study were surprisingly targeted to *fimH*. To test the evolution towards increased adhesion capacity in absence of *fimH*, we deleted the whole *fim* operon (*fimABCDEFGH* see Supp. Fig. S3A) and we ran 12 parallel positive selection experiments of the *Δfim* strain for 15 cycles. We observed an evolution towards significant and relevant increased biofilm formation in both biofilm models for 3 out of 12 *Δfim* populations, ^Δ^G2, ^Δ^G5 and ^Δ^R4 (Fig. 6A-B), with ^Δ^G5 and ^Δ^R4 achieving 14- and 71-times increased biofilm capacity in biofilm microfermenters (Fig. 6A). The analysis of the mutations found in these three biofilm-positive evolved *Δfim* populations and in corresponding clones revealed mutations affecting biofilm associated functions such as chaperone-usher fimbrial surface structure (*yqiG*, *ecpR* and *ecpD*), autotransporter adhesin (*ycgV*, *flu*/*agn43*) and flagellum (*flgH*, *ecpR*) (Supp. Table S6).

**Figure 6.**
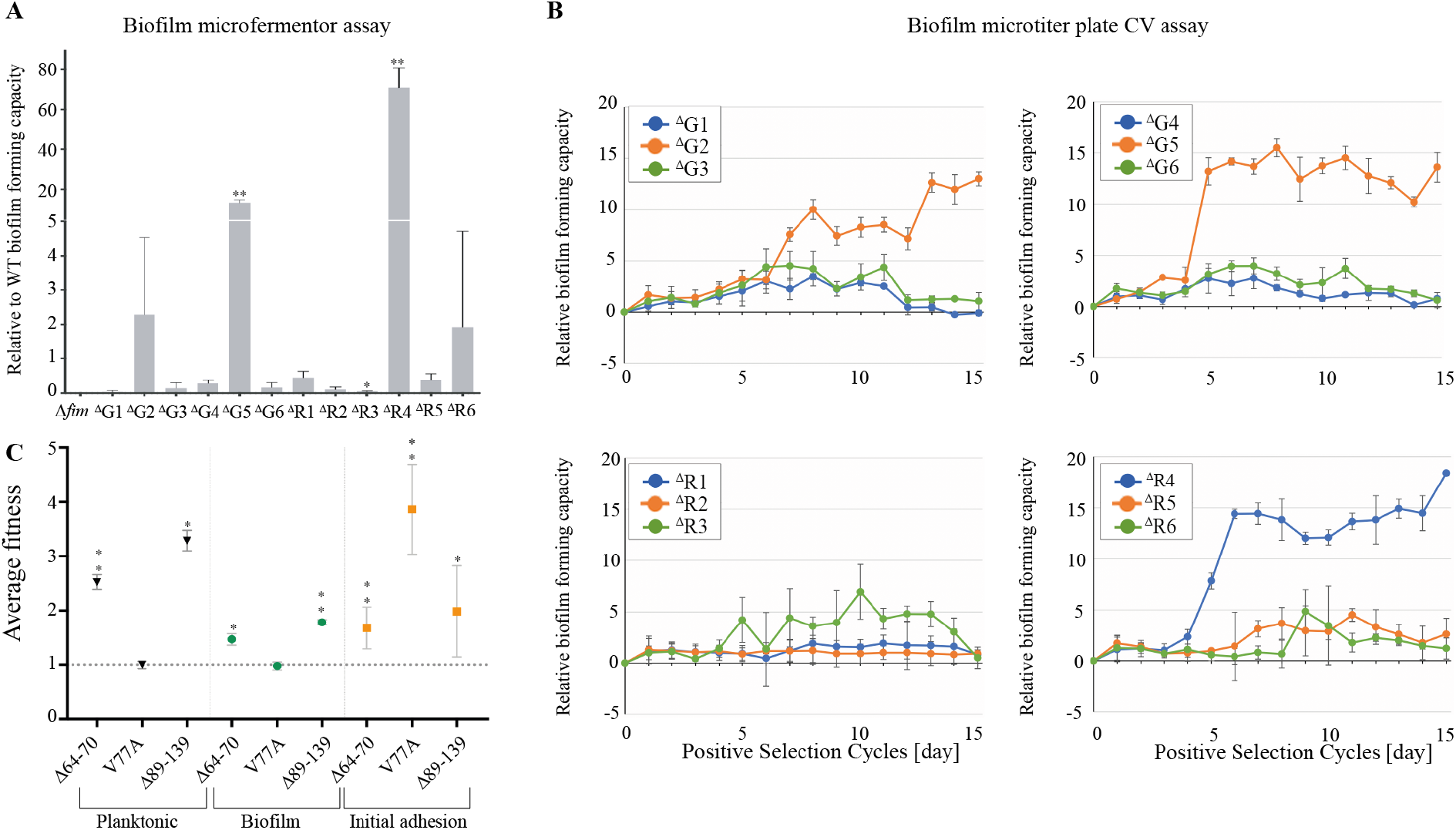
Evolution of biofilm forming capacity of *Δfim* bacterial populations subjected to positive selection for adhesion. **A.** The relative biofilm forming capacities of the populations at the end of the ^Δfim^positive selection experiment, after 24 hours in microfermenters. The relative capacities are calculated using the capacities of the wild type (1) and *Δfim* ancestral strains (0). All microfermenter biofilm experiments were performed in triplicate. to unpaired t-test with Welch’s corrections. **B.** The biofilm forming capacities were monitored at each cycle during the ^Δfim^positive selection experiment with the crystal violet assay of an aliquot of the enriched population as well as the parental wild type strains. The relative capacities were calculated using WT *E. coli* MG1655 *<u>Δfim</u>* (very poor biofilm former) and *E. coli* MG1655 strain as follows: (OD_test_-OD_Δfim_)/(OD_MG1655_-OD_Δfim_).. When the value is equal to 1 and 0, the capacity is as same as the WT parental and the Δ*fim* strains, respectively. Each panel shows three populations: ^Δfim^G1-3, ^Δfim^G4-6, ^Δfim^R1-3 and ^Δfim^R4-6. All crystal violet biofilm assay experiments were conducted with at least 8 replicates. **C.** Comparison of fitness for three *fimH* mutants under native promoter competed against the best biofilm former mutant from the Δ*fim* evolution experiment (ecpR IS2(-) +5 bp / ecpD G89D). The strains were competed for planktonic growth, biofilm formation and initial adhesion on the microfermentor glass spatula as described in Materials and Methods. Dotted line represents *ecpD/ecpR* mutant fitness set to 1. Statistics correspond to unpaired, non-parametric Mann-Whitney test comparing all conditions to WT *Δfim*. * p<0.05; ** p<0.01.

Whereas the *ecp* operon is not expressed in *E. coli* K12 MG1655 ^31^, biofilm-promoting mutations in *ecpD* were only found in clones also displaying an IS insertion right after *ecpR*, the regulator of the operon. This IS insertion resulted in *ecp* operon expression (Supp. Fig. S9), potentiating further mutations in *ecpD*, leading to increased biofilm formation in *Δfim* population ^Δ^R4. These results therefore showed that, in absence of type 1 fimbriae, different mutations in surface structure and adhesins could lead to increased biofilm capacity.

### *fimH* mutants outcompete mutants selected in the absence of *fimH*

To investigate the potential origin of the lack of mutation diversity in evolution experiments performed in the WT background compared to those performed in a Δ*fim* background, we reconstructed the *ecpD/ecpR* mutant from the ^Δ^R4 population (the highest biofilm-forming population from the Δ*fim* evolution) in a WT background. This mutant was then used in competition assays against the natural *fimH* mutants with highest biofilm capacity (Δ64-70 and Δ89-139) and increased, but lower biofilm capacity (V77A). These competition experiments were performed for three critical steps of our selection protocol: initial adhesion on the glass spatula inserted in biofilm microfermenters, biofilm formation and growth in liquid culture. We showed that, during biofilm formation in microtiter plates as well as growth in liquid culture, both Δ64-70 and Δ89-139 *fimH* mutants outcompeted the *ecpD/ecpR* mutant (Fig. 6 C). Whereas there was no advantage of the V77A mutation in these conditions, competition for initial adhesion showed a very clear advantage for the V77A *fimH* mutant and, in a lesser extent, for both Δ64-70 and Δ89-139 mutants (Fig. 6 C). These observed fitness advantages of the *fimH* mutants could account for the out-competition of other adhesin mutations in our evolution experiments.

## DISCUSSION

In this study, we showed that despite the vast arsenal of proteinaceous and macromolecular surface structures known to contribute to biofilm formation in *Escherichia coli*, *in vitro* experimental evolution selecting for increased biofilm-forming capacities systematically led to the acquisition of mutations in the type 1 fimbriae tip adhesin gene *fimH*, one of the first *E. coli* appendages implicated in biofilm formation on abiotic surfaces ^21^. Although the restricted mutational landscape revealed by our study suggests that type 1 fimbriae are the main *E. coli* adhesins in these experimental conditions, we cannot exclude that mutations in other genes could have emerged and been counterselected before the last selection cycles or been present below the 5% threshold detection of our *breseq* analyses. For instance, the G2 population showed an episodic increase in adhesion capacity at day 11 that was not detected afterwards (Fig. 1B). This correlated with the presence of an in-frame deletion in *flu*, the gene coding for the self-recognition *E. coli* adhesin Antigen 43 ^32, 33^. We hypothesize that this mutation and other potential *fimH*-independent mutants could be outcompeted by emerging *fimH* mutations leading to a strong bias toward type 1 fimbriae mutations. In support of this hypothesis, we showed that, although positive selection for increased adhesion in a strain deleted for the *fimA-H* operon also identified mutations in genes encoding adhesins, the strongest biofilm-former amongst these mutants was outcompeted by *fimH* mutants for growth, initial adhesion and biofilm formation. In addition, we observed that mutation of the tip-pilus adhesin *ecpD* was always observed together with the insertion of an IS element that increases expression of the otherwise cryptic *ecp* pilus operon. This suggests that adhesin expression may first need to be unlocked before productive mutations could enhanced their adhesive properties and our selection could favor bacteria in which *fim* expression was locked ON, so more likely to evolve towards increased adhesion.

FimH is an allosterically regulated mannose-binding protein and FimH-dependent mannose-binding is considered as important for adhesion to mannosylated cell surface receptors *in vivo* as well as *in vitro* biofilm maturation possibly via the recognition of mannose-rich biofilm matrix component ^34^. In presence of shear forces, FimH mannose binding-capacity is enhanced by catch-bond mechanisms ^25, 35^, which could provide a selective advantage in the turbulent conditions used to select for increased-adhesion mutants. However, while most identified *fimH* mutants displayed an enhanced capacity for initial adhesion to abiotic surface they exhibited reduced mannose-binding capacity and *fimH* mutants with deletion encompassing part of FimH mannose-binding pocket were amongst the strongest biofilm-formers. This shows that FimH-dependent mannose-binding and catch bond do not significantly contribute to the increased biofilm capacities of the mutants identified in our study. FimH was early demonstrated to contribute to both specific and non-specific adhesion ^21^ and our results show that the lectin domain of FimH contributes to both types of adhesion, with residues of this lectin domain directly engaged in interaction with abiotic surfaces. This revealed a trade-off between attachment to mannose and abiotic surfaces that could be due to the fact that residues that are mutated or deleted in our selected *fimH* mutants are either at the vicinity of the mannose binding pocket and could directly impact the specificity of the interaction through structural modification (Supplementary Fig. S10). Hydrophobic interactions also play a key role in bacterial adhesion ^36^ and mutations reducing hydrophobicity could enhance FimH interactions to abiotic surfaces. Such mutations were selected during our evolution experiments: substitution of hydrophobic by hydrophilic amino acid residue (V77A, L79P), producing the mutants with the highest initial adhesion on the spatula (Fig. 5), substitution of non-polar by polar amino-acid (G82C and G87C) or deletion of stretches of hydrophobic residues (Δ89-139 correspond to the deletion of 38 hydrophobic residues out of 51). Hence, increasing FimH hydrophilicity could contribute to improve adhesion of FimH mutants on microfermenter glass spatula. In the case of the S16L mutant in FimH signal peptide, alteration of signal peptide could increase FimH transport efficiency and type 1 fimbriae exposition ^37^. However, we could not detect any differences in the quantity of surface exposed FimA in this FimH S16L mutant. Alternatively, the selected mutations in *fimH* could improve surface contact and adhesion due to changes in the tertiary structure of FimH (Supplementary Fig. S10).

Unexpectedly, our study also revealed a negative correlation between the strength of initial adhesion displayed by the evolved *fimH* mutants and their capacity to form mature biofilms. This trade-off is particularly clear in the case of mutations at position V77A and L79P, leading to strong increase in initial adhesion with almost no positive impact on biofilm formation capacity. By contrast, the Δ64-70 and Δ89-139 mutations showed low gain in initial adhesion but led to strong increase in biofilm formation. These results illustrate the complexity of the mechanisms at play during biofilm formation. Strong initial adhesion could deeply impact bacterial metabolism and significantly delay biofilm maturation ^38^. Alternatively, but these hypotheses are not mutually exclusive, a too strong attachment to the surface might not be optimal to favor later cell-to-cell interaction and matrix production, but also the dynamism, including movement of cells, that might be necessary during biofilm maturation.

Type 1 fimbriae were shown to contribute to both pathogenic and commensal *E. coli* colonization of biotic surfaces. FimH-mediated adhesion enables commensal *E. coli* to adhere to buccal and intestinal epithelia as part of the normal bacterial flora but also enables pathogenic strains to colonize various mannosylated tissues ^39–41^. Although mutations in the FimH pilin domain are likely negatively selected because of its critical functional role, the mannose lectin domain displays a higher genetic plasticity, which could lead to functional diversity, conferring selective advantage by modifying and diversifying substrate binding capacity ^42, 43^. Consistently, several phenotypic variants of the FimH lectin domain have been identified in clinical urinary or intestinal *E. coli* isolates ^28, 44–48^. These variants are often single point, non-synonymous amino acid substitutions found in the lectin domain but not affecting the mannose-binding pocket directly. While adaptive mutations identified in our study were also found in the lectin domain of the protein, only position 87 (G87C) has been previously described in pathoadaptive variants ^28, 44, 45, 48^ and described as a mutational hotspot with amino acid changes to arginine, alanine, serine and cysteine ^28^. The G87S and G87C variants are the only mutants that display moderate increases in mannose-binding, but still display catch-bond properties under flow conditions ^28^.

While FimH variants previously identified in clinical isolates were mostly associated to tissue tropism ^28, 45, 46, 48^, our study shows that some mutations isolated in natural and clinical strains can also be selected for increased initial adhesion to abiotic surface and, as a consequence, biofilm formation capacity. This suggests that evolution of FimH in natural and/or clinical environments can also be driven by abiotic environmental selection pressures. The observation that FimH sequence distribution does not strictly reflect phylogroup clustering (Supp. Fig. S7) further suggests that this gene is subjected to selection pressures specific to each strain environment and, consequently, the need for specific adaptations. Supporting this idea, the mutational landscape analysis of FimH sequences in *E. coli* revealed 20 hotspots constituting a potential source of functional diversity. These different hotspots have various phylogroup enrichment suggesting that these mutations are related to diverse ecological contexts and environments. For instance, the co-occurrence of S91N and N99S related to UPEC strains ^27^ was also identified in environmental strains, which is consistent with the growing evidence supporting the cycling of *E. coli* between the host gut and the environment^49^. Although the frequency of this lifestyle switch and how long each *E. coli* clone resides in the gut and in the environment is not well known, *E. coli* is able to grow, evolve and adapt in both of these environments. Interestingly, conditions in the mammalian gut are relatively stable as compared to the one found in the outside host environment, and experimental evolution of *E. coli* in the gut did not select for mutation in *fimH* ^50–54^, in contrast with studies on plant-associated *E. coli* demonstrating an evolution towards increased capacity to form biofilm in this environment ^55^. Hence, rapidly changing conditions encountered by *E. coli* outside of the host could impose strong selection pressures on environmental *E. coli*, potentially leading to their rapid evolution ^56, 57^. The fact that FimH mutations selected in our experiment correspond to naturally occuring mutations enriched in B2 and F phylogroups (mainly ExPEC strains), suggests that mutations favouring non-specific *versus* specific adhesion could be selected even in strains known for their tissue tropism. One can therefore speculate about the nature of the main *in vivo* and/or environmental selection pressure driving the evolution sequenced *E. coli* clones. Unfortunately, the clear lack of environmental *E. coli* strains in the databases hinders genomic analysis. Indeed, among the 3543 FimH sequences included in our analysis 1256 were of unknown origin, 1192 were isolated from human, 690 from animals, 64 from food-related samples and only 341 from the environment. This therefore emphasizes the need for an increased number of sequenced and annotated genomes of environmental *E. coli* strains.

From an evolutionary perspective, the benefits of FimH mutations affecting mannose binding capacity greatly depend on the environmental surfaces colonized by *E. coli*. FimH variants with increased mannose affinity under shear stress could be advantageous for bladder colonization, while decreased concentrations of free soluble mannose in the intestine could impose a very different selection pressure. Our findings show that *fimH* can undergo rapid microevolution, leading to increased non-specific adhesion and biofilm formation independently of mannose-binding capacity. This extended FimH mutational landscape in biofilms could reflect a natural strategy to diversify *E. coli* surface binding capacity, resist physical and chemical disruptions and potentially providing selective advantage for persistence during *E. coli* cycling between hosts and the environment.

## MATERIAL AND METHODS

### Bacterial strains and growth conditions

Bacterial strains used in this study are listed in supplementary Table S7. *E. coli* was grown in M63B1 minimum medium supplemented with glucose (0.4 %) and kanamycin (20 µg/mL), and incubated at 37°C. All media and chemicals were purchased from Sigma-Aldrich.

### Strain construction

The WT strains used in the experiments originated from an *E. coli* MG1655 K12 wild-type strain and were either tagged, at the lambda att site, with red (*mars*) or green fluorescent protein encoding genes (*gfpmut3)*(RFP/GFP) by P1 vir transduction. Strains without the whole *fim* operon (Δ*fimABCDEFGH*) were constructed by P1 vir phage transduction method from MG1655_Δ*fimAICDFGH::cat* ^6^ into the WT strains. The constitutive promoter controlling the *fim* operon (PcL*fim*) ^7^ was transferred by lambda-red recombination into WT strains and individual *fim* mutants using pKOBEGA plasmid and lambda red recombination. The *ecpD/ecpR* mutant was reconstructed using P1 vir phage transduction into the MG1655 K12 wild-type strain tagged with either red (*mars*) or green fluorescent protein (*gfpmut3)*. Primers used for genetic contruction are listed in supplementary Table S8.

### Biofilm formation in microfermenters

Continuous-flow biofilm microfermenters containing a removable glass spatula were used as described in ^58^, with internal agitation provided by filter-sterilized air-bubbling. Biofilm microfermenters were inoculated by placing the spatula in a culture solution adjusted to OD_600_=1.0 (containing 5.0x10^8^ bacteria/mL) for 10 min. The spatula was then reintroduced into the microfermenter and biofilm culture was performed at 37°C in M63B1 with 0.4% glucose. Flow rate was then adjusted (30 mL/h) so that total time for renewal of microfermenter medium was shorter than bacterial generation time, thus minimizing planktonic growth by constant dilution of non-biofilm bacteria.

### Positive selection procedure

At the beginning of the positive selection procedure (Day 0) -80°C glycerol stocks of the parental *E. coli* GFP- or RFP-tagged strains were inoculated in culture tubes containing LB medium supplemented with kanamycin (20 µg/mL) for over-day 8h culture, then transferred into M63B1 minimum medium supplemented with 0.4% glucose and kanamycin for overnight 15 h culture. Six removable glass spatula were submerged into the GFP- overnight culture adjusted to OD_600_ = 1.0 (ca. 5.0x10^8^ bacteria/mL) for 10 min, and six others with the RFP- overnight culture. Each spatula was then reintroduced into one microfermenter and the bacteria adhered to the spatula were incubated for eight hours, during which fresh medium (M63B1 minimum medium supplemented with 0.4% glucose) flowed (30 mL/h) in the microfermenter, constantly diluting non-adhering, planktonic bacteria. After the eight hour incubation in the microfermenters, bacteria that developed as a biofilm onto the spatula were resuspended in culture tubes containing fresh minimum growth medium supplemented with glucose and kanamycin, and cultured overnight. A fraction of each overnight culture was used to prepare a glycerol stock that could be analyzed afterwards and the rest was then used for the inoculation for the next cycle of the positive selection experiment.

Microfermenters were sterilized using 70% ethanol overnight after each positive selection cycle, then air dried prior to the following positive selection cycle. The sterility of the microfermenters was tested by plating out an aliquot of the fresh medium added to the microfermenters at the beginning of each positive selection cycle. The spatula was autoclave-sterilized, machine-washed, then sterilized again between the cycles. During the 15 days of the positive selection procedure, aliquots of the population were stocked at -80°C (glycerol stock) and the adhesion capacity of the populations was monitored by running crystal violet microtiter plate biofilm assay. At the end of the positive selection cycles, bacteria were resuspended from the spatula and these selected end-point populations were stocked. The biofilm formation capacity of these end-point populations were compared to those of non-evolved parental wild type GFP- or RFP-tagged strain after 24 h in microfermenters by monitoring microfermenters and spatula and biomass after resuspension in M63B1 (no glucose) and determination of OD_600_ optical density. These 24 h microfermenter experiments were performed at least three times.

### Biofilm microtiter plate assay CV assay

The glycerol stocks of the target strains or populations were inoculated in LB medium supplemented with kanamycin (20 µg/mL) for over-day 5h culture at 37°C, then inoculated in either a polystyrene Greiner 96-well plate or a polyvinyl chloride Corning plate at OD_600_ =0.01 (5x10^6^ cells/mL) in M63B1 minimum medium supplemented with 0.4% glucose and kanamycin. Incubation was done at 37°C for overnight, 16 hours. The supernatant was removed from each well and Bouin solution (Sigma) was applied for 15 minutes to fix the biofilm attached to the well. Next the fixation solution was washed 3 times with water and biofilm was stained with 1% crystal violet solution (QCA) for 15 minutes. After removal of the crystal violet solution, biofilms were washed 3 times with water. For quantification of biofilm formation, dried stained biofilms were resuspended in 30% acetic acid and absorbance was measured at 585 nm using an infinite M200 PRO plate reader. Biofilm formation in different strains is represented in values normalized to average biofilm formation in the control strains.

### Competition assay for biofilm formation

The glycerol stocks of the target strains were inoculated in LB medium supplemented with kanamycin (20 µg/mL) for over-day 6h culture at 37°C, then overnight in M63B1 minimum medium supplemented with 0.4% glucose and kanamycin. After overnight incubation, the cultures were adjusted to OD_600_ = 1.0 (ca. 5.0x10^8^ bacteria/mL). The target strains with different fluorescent tags were mixed in 1:1 ratio and the cell concentrations were verified by FACS and OD analyses. Each competition mix was then inoculated in either a polystyrene Greiner 96-well plate or a polyvinyl chloride Corning plate at OD_600_ = 0.01 (5x10^6^ cells/mL) in M63B1 minimum medium supplemented with 0.4% glucose and kanamycin. Incubation was done at 37°C overnight, for 16 hours. The supernatant was removed from each well and the biofilm was resuspended in 100 µL of PBS. The proportion of each strain was then assessed from this resuspension by FACS and the relative fitness of target strain was calculated as follows:

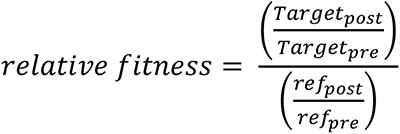

where *Target*_*pre*_ and *Target*_*post*_ are the cell concentrations of the target strain in the mixed culture before and after the overnight incubation, and *ref*_*pre*_ and *ref*_*post*_ are the cell concentrations of the reference strain in the mixed culture before and after the overnight incubation, respectively ^59^.

### Competition assay for growth in liquid cultures

The glycerol stocks of the target strains were inoculated in LB medium supplemented with kanamycin (20 µg/mL) for over-day 6 h culture, then for overnight in M63B1 minimum medium supplemented with 0.4% glucose and kanamycin. After overnight incubation, the cultures were adjusted to OD_600_ = 1.0 (ca. 5.0x10^8^ bacteria/mL). The target strains with the different fluorescent tags were mixed in 1:1 ratio and the cell concentrations were verified by FACS and OD analyses. Then they were transferred into M63B1 minimum medium supplemented with 0.4% glucose and kanamycin at 1000 times dilution and incubated overnight. The concentrations of target and reference strains in the overnight culture was measured using FACS and spectrophotometer. The relative planktonic fitness during overnight culture were calculated as for competitions for biofilm formation assay (see above).

### Competition assay for initial adhesion

Prior to the initial adhesion assay, the glycerol stocks of the target strains were inoculated in culture tubes containing LB medium supplemented with kanamycin (20 µg/mL) for over-day 6h culture, then transferred into M63B1 minimum medium supplemented with 0.4% glucose and kanamycin for overnight culture at 37°C. The overnight culture were adjusted to OD_600_ =1.0 (ca. 5.0x10^8^ bacteria/mL) and the target and reference strains with the different fluorescent tag were mixed in 1:1 ratio. The ratio was verified by FACS analysis. Glass spatulas were inoculated with 15 mL of the mixed culture for 10 mins and washed 3 times in 20 mL PBS. The bacteria attached to the glass spatula were resuspended in 10 mL minimum medium without glucose. The ratio of each strain in the resuspended solution was measured using FACS. The results were either expressed as (i) relative fitness for Figure S2 and were calculated as for competitions for biofilm formation assay (see above), or (ii) for Figure 4 as percent gain within the population as follows;

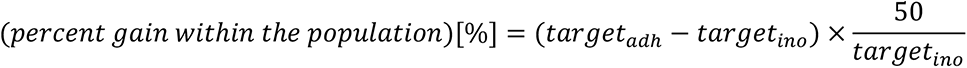

where the measured percentage of the target strain in the inoculation mixture and the resuspention are shown as *MT_ino_* and *MT_adh_*, respectively.

### RNA extraction and RT-PCR

Three colonies of each strain were grown in LB medium supplemented with kanamycin (20 µg/mL) for over-day 6h culture at 37°C, then overnight in 25 mL of M63B1 minimum medium supplemented with 0.4% glucose and kanamycin. RNA extractions were then performed using Trizol Reagent (Ambion) following manufacturer instructions. Briefly, the equivalent of 5.0x10^7^ cells were centrifuged for 5 minutes at 5,000 rpm, the pellets were resuspended in 1 mL of Trizol Reagent and incubated 5 minutes at room temperature. 200 µL of chloroform was added and each tube was vortexed for 15 seconds and incubated for 5 minutes at room temperature before centrifugation for 15 minutes at 10,000 rpm and at 4°C. The upper phase containing the total RNA was then transferred in 500 µL of isopropanol. The tubes were mixed by inversion and incubated for 5 minutes at room temperature before centrifugation for 10 minutes at 10,000 rpm and at 4°C. The supernatant was discarded and the pellets were washed by adding 1 mL of 70% ethanol. The tubes were mixed by inversion and incubated for 5 minutes at room temperature before centrifugation for 10 minutes at 10,000 rpm and at 4°C. After removing the supernatant, the pellets were air dried and resuspended in 50 µL of RNAse free water. A DNAse treatment was then applied to all samples using the TURBO DNAse kit (Ambion). The resulting RNA was used as a matrix for reverse transcription using the AMV cDNA first strand synthesis kit (Roche). The produced cDNA was used to assess *ecpD* expression as well as 16S rRNA gene as a control.

### Whole genome sequencing and analysis

Prior to genome extraction, the glycerol stocks of the target populations and clones were inoculated in LB medium for over-day culture till the OD_600_ reached around 1.0 (ca. 5.0x10^8^ bacteria/mL). The bacterial cells were collected from 2 mL of the culture and the genomic DNA was extracted using the Qiagen DNeasy Blood and Tissue kit. RNase digestion step was added during the genome extraction process. Sequencing libraries were prepared using Nextera XT or Nextera Flex DNA Library Preparation Kit. All samples were sequenced using Illumina HiSeq sequencer. Sequencing reads were pre-processed to remove low-quality or artefactual bases. We used fqCleaner v.0.5.0, a mini workflow implemented in Galaxy ^60^ to process fastq files (quality trimming, duplicate and artifact filters). Mutations with a frequency superior to 5% were detected using *breseq* version 0.30.0 ^61^ with the consensus mode for clones sequences analyses and the polymorphism mode for population sequencing. In both cases default parameters were used.

### Determination of the frequency of *fimH* mutation

The evolution of frequency of the identified mutations in *fimH* was performed using Sanger sequencing analysis of PCR products centered on the *fimH* region using the following oligonucleotides - oligo up: AGGATGACAGTGGCAACACA and oligo down: GTTTTGGCTTTTCGCACAAT. Small aliquots of the glycerol stocks corresponding to each positive selection cycle were diluted in water and used for PCR reaction (Thermo Phusion flash high-fidelity master mix). The PCR products were sent to Eurofins for purification and Sanger sequencing. The frequency of the mutations was calculated using QSVanalyzer ^62^, with a cutoff that does not allow detection of mutations lower than 5% frequency.

### Determination of *fim* “ON/OFF” status of evolved clones

Orientation of the 314 bp DNA segment harboring the *fimA* promoter (*fimS* region) was determined using a PCR-based assay using restriction fragment length dimorphism arising from the orientation-dependent location of a unique BstUI restriction site within the amplified DNA. Briefly, the switch (*fimS*) region was amplified from a sample of overnight cultures with oligonucleotides OL4 (5’ CCGTAACGCAGACTCATCCTC 3’) and OL20 (5’ GAGTTTTAATTTTCATGCTGCTTTCC 3’) to generate a 726 bp PCR product. DNA was amplified with Taq polymerase (Invitrogen) using the following PCR conditions: denaturing at 94°C for 5 min, followed by 30 cycles (94°C for 1 min, 58°C for 1 min and 72°C for 1 min) and a final extension of 10 min at 72°C. Samples were cooled at 8°C, 10 units of BstUI (Nex England Biolabs) were added to each reaction and incubation was conducted at 37°C for 3 h. Digested PCR products were resolved on 2% agarose gels. Using this assay, phase ON populations of bacteria yielded two DNA fragments 433 and 293 bp in length, whereas phase OFF populations yielded two fragments of 539 and 187 bp. Mixed populations contained a mixture of all four fragments.

### Protein modelling and structure prediction

The 3D structure of the lectin domain of MG1655 FimH presented corresponds to the pdb model 1KLF ^63^. The 3D structures of the mutant FimH proteins were predicted using the PHYRE2 Protein Fold Recognition Server and visualized with the MacPymol Software ^64, 65^.

### Type 1 fimbriae extraction

Surface exposed type 1 fimbriae was isolated by heat shock extraction: a 5 mL culture of each strain was grown in LB at 37 °C for 16 h and OD_600_=10.0 equivalent of cell culture was harvested by centrifugation. The harvested cells were washed with 0.9% NaCl, collected by centrifugation and resuspended with 75mM NaCl, 0.5mM Tris-HCl, pH 7.4. The samples were incubated at 60°C for 20 min, cooled on ice for 3 min, centrifuged, and then the detached adhesins present in the supernatant were precipitated with 10% TCA overnight. The proteins were collected by centrifugation (20,000 g, 1 h, 4°C), then the acquired pellet was washed with 75% acetone and the proteins were dissolved in HEPES 10 mM.

### Detection of FimA by Western blot with anti-FimA antibodies

The heat extracted proteins from OD_600_=2 culture was suspended in 1X Laemmli buffer with 250 U of benzonase nuclease (Sigma E0114) and incubated for 5 minutes at 95°C. The protein extracts were run on Mini-PROTEAN TGX Stain-Free^TM^ precast Gels (BioRad) in 1X TGX buffer and then transferred to nitrocellulose membrane using a Trans-Blot^®^ Turbo™ Transfer System (BioRad). Blocking was performed in a 5% solution of dry milk and 0.05% Tween –1X PBS (1X PBST) overnight at 4 °C with agitation. The membranes were then incubated in 1X PBST with a polyclonal rabbit antiserum raised against FimA subunit of type 1 fimbriae (kindly given by Prof. Scott Hultgren) at 1:10000 for 1 h at room temperature with agitation. Membranes were washed in 1X PBST and then incubated with the secondary antibody (anti-rabbit IgG conjugated with horse radish peroxidase at 1:10000, Promega). After washing the excess secondary antibody, specific bands were visualized using the ECL prime detection method (GE Healthcare).

### Yeast agglutination assay

The capacity of WT and *E. coli* mutants expressing FimH variant to bind yeast mannosylated proteins and agglutinate yeast cells was assessed as previously described ^66^. The glycerol stocks of the PcL mutants and the PcL WT strains were inoculated in culture tubes containing LB medium supplemented with kanamycin (20 µg/mL) for over-day 8h culture, then transferred into M63B1 minimum medium supplemented with 0.4% glucose and kanamycin for overnight culture. The culture were washed once with 20 mL PBS and resuspended at OD_600_ = 9 in PBS. 2 % w/v yeast suspension was prepared using 100 mg dry *Saccharomyces cerevisiae* (Sigma) and 5 mL PBS. A range of 12 two-fold dilutions of bacterial cells from OD600 = 9 to OD600 = 0.02 was tested for each strain. 50 μL serial dilutions of bacteria in PBS were mixed with 50 μL yeast suspension in round-bottom 96 well-plates (Fisher). After 3 hours of incubation (static condition) or 30 mins with 600 rpm shaking (dynamic condition) at room temperature, the lack of agglutination was seen as an aggregation of yeast cells at the bottom of the well. In the presence of agglutination there was no yeast clumps but a homogeneous mixture of yeast and bacteria. The agglutination titer corresponding to the lowest concentration of bacteria leading to yeast agglutination was recorded for each strain.

### FimH sequences analysis

The FimH sequence from the ancestral strain (*E. coli* K12, strain MG1655) used in the positive selection experiment was used as a query for BLASTp searches (blast+ version 2.2.31, ^67^ against the NCBI nrprot database (version 2019-04-09) as well as against a custom database composed of 2067 sequenced genomes of *E. coli* (supplementary Table S2) and 277 genomes from environmental strains published in ^26^. Only hits corresponding to *E.coli* and with an e-value lower than 1e^-10^ and sequence identity higher than 70% were kept. In our analysis, the detection of one mutation in two identical sequences coming from different origin was important. Therefore, the redundancy of the FimH sequences dataset was not addressed in terms of sequence identity, but rather in term of isolates. To avoid such duplicated sequences, FimH proteins with identical identifiers or strain information were discarded. The resulting non-redundant dataset was composed of 3543 sequences (supplementary Table S3AB).

All sequences were aligned using mafft version 7.407 ^68^ with G-INS-i option and the resulting alignment was screened to identify mutations relative to the reference sequence from *E. coli* K12, strain MG1655 using custom perl script. The same alignment was used to compute the single amino-acids modifications frequencies for each sequence in the signal peptide, mannose lectin domain, linker part and the pilin domain. For each sequence, the mutation frequency was computed for the whole sequence, the signal peptide, the lectin domain and the pilin domain. These dataset were then compared using a one-way Anova. To identify positions with a significantly higher mutation frequency, the same method was perfored using a sliding winow approach.

### Selection pressure analysis

The *ω* ratio of non-synonymous (d_N_) to synonymous (d_S_) nucleotide substitutions is widely used to estimate the evolutionary forces acting on a gene ^69^. A ratio of 1 is considered to indicate neutral evolution while a ratio higher than one indicates positive selection and a ratio lower than one indicates purifying selection. PAML version 4.9 ^29^ was used for the positive selection analyses. First, only unique sequences (100% identity) were kept in the previous alignment (533 sequences), which was trimmed using trimal v1.4.1 ^70^ and used as a guide to codon align the related nucleotide sequences using PAL2NAL version 14 ^71^. The resulting alignment was then used to compute a phylogenetic tree using PhyML version 3.1 ^72^ with 100 bootstraps and both were used as input for PAML. The best fit model for phylogenetic tree reconstruction was inferred using jModelTest2 version 2.1.3 ^73^. The phylogenetic tree of Supp. Figure S8 was visualized and edited using iTOL v4 ^74^.

The site models M0, M3, M8a and M8 were first run on the whole tree. The comparison of M0 vs M3 tests for rate heterogeneity between amino acid sites, while M8a vs M8 tests for positive selection. These tests only detect positive selection acting on a site when the *ω* ratio averaged over all branches is higher than one. However, positive selection could be restricted to a sub-group of sequences, which is diluted in the whole tree. To identify such sub-groups of FimH sequences subjected to positive selection we used two different strategies. (i) We first divided the tree in four sub-groups (Sg1 to Sg4) based on visual inspection and phylogroup distribution along the tree as well as positions 70 and 78 (91 and 99 in this study) for which alleles S and N, respectively, were described to be specific to UPEC, which allows the discrimination of Sg4 and Sg3. The remaining sequences that were not assigned to Sg1 to Sg4 were put in a fifth sub-group. Then we used the branch-site model A, which allows *ω* to vary among sites in a specific branch of the tree (the foreground branch) ^75^, by labeling Sg1 - Sg4 as the foreground branch. Since Sg5 is not a monophyletic group, it was not investigated using this method. (ii) The second strategy was to build new alignments and trees for each of the five sub-groups and run models M0, M3, M8a and M8 on each of them separately.

Finally, to test if the lectin domain and the pilin domain evolved at different rates and were subjected to different selective pressures, only these two domains were kept in the different alignments (whole tree and sub-groups 1 to 5) and labeled as two different site partitions. Five different fixed-site models were then run ^76^: model A assuming a single *ω* ratio for the entire sequence; model B assuming different substitution rates; model C assuming different substitution rates and codon frequencies; model D assuming different substitution rates, different transition/transversion ratios and different *ω* ratios among the partitions; and model D2 which is the same as model D but with a fixed transition/transversion ratio. The comparison of model A vs model B tests for evolution of the two partitions at different rates. The comparison of model B vs model C tests for different codon usage between the two partitions. The comparison of model B vs model D tests for different transition/transversion and *ω* ratios between both partitions. The last comparison between model B and model D2 allows to test for different *ω* ratios only.

In each case, a likelihood ratio test assuming a *χ*2 distribution was used to compare each pair of models using twice the difference in log likelihoods as *χ*2 and the parameters difference reported by PAML as degrees of freedom. The resulting p-values were corrected in order to account for multiple testing and a FDR threshold of 0.05 was considered as significant. For such statistical tests, posterior probabilities under Bayes Empirical Bayes ^77^ analysis under M8 model and branch-site model A were extracted to identify sites under positive selection. To avoid local optima, all analyses were run with different starting *ω* values (0.04, 0.4 and 4).

### Statistical analysis

Unpaired, non-parametric Mann-Whitney tests were performed using Prism 6.0 for Mac OS X (GraphPad Software, Inc.) for CV staining biofilm assay in which each experiment was performed at least 8 to 12 times. Unpaired t-test with Welch’s corrections were performed in the case of continuous flow biofilm experiments in which each experiment was performed 3 times.

## DATA AVAILABILITY

All sequencing reads were deposited in NCBI under the BioProject accession number PRJNA714528. The perl scripts used are available at https://github.com/Sthiriet-rupert/FimH_Evolution

## ACKNOWLEDGEMENTS.

We thank Rebecca Stevick, Médéric Diard and Olaya Rendueles for critical reading of the manuscript. We are grateful to Prof. Scott Hultgren for kindly providing the anti-FimA antiserum and to Dr. Olaya Rendueles for the initial help with the analysis of the mutations. This work was supported by an Institut Pasteur grant, by the French government’s Investissement d’Avenir Program, Laboratoire d’Excellence "Integrative Biology of Emerging Infectious Diseases" (grant n°ANR-10-LABX-62-IBEID) and by the Fondation pour la Recherche Médicale (grant DEQ20180339185). M.Y. was supported by Institut Pasteur Roux Cantarini fellowship. S.T.-R was supported by the French National Research Agency (ANR), project EvolTolAB (ANR-18-CE13-0010).

## AUTHOR CONTRIBUTIONS

M.Y., C.B. and J.-M.G. designed the experiments. M.Y., L.M. and S.T.-R. performed the experiments. M.Y., C.B., S.T.-R., L.M. and J.-M.G. analyzed data. J.-M.G, C.B., M.Y. and S.T.-R. wrote the paper.

## SUPPLEMENTARY MATERIAL

### SUPPLEMENTARY TABLES

Supplementary Table S1(available https://research.pasteur.fr/en/project/supplementary-tables-for-biorxiv/): **Mutations identified in populations and clones from the evolution in a WT genetic background at cycle 15 of the experiments**. Datasheet A in the 6 biofilm positive populations, Datasheet B in the 6 biofilm negative populations. For each mutation identified the gene, nature of the mutation, impact on the protein sequence as well as the frequency of the mutation in the population are shown.

Supplementary Table S2 (available https://research.pasteur.fr/en/project/supplementary-tables-for-biorxiv/): ***E. coli* genomes used in this study** For each genome, the strain’s NCBI taxonomy identifier is shown in the first column, the name of the corresponding strain in the second column and the related NCBI genome assembly identifier in the third column.

Supplementary Table S3 (available https://research.pasteur.fr/en/project/supplementary-tables-for-biorxiv/): **Summary table of all FimH sequences used in this study.** For each sequence, the effective length is specified for the signal sequence, the mannose lectin domain, the linker part and the pilin domain as well as for the whole protein. The number of mutations and the related mutation frequencies (number of mutations per amino acid) are also specified. Features obtained from NCBI (Strain name, serotype, isolation source and host) are noted when available. The phylogroup was determined using ClermonTyping method (Beghain et al., 2018) and positions 70 and 78 (numbered 91 and 99 in this study) that were described as related to UPEC strains are also specified. Finally, sequences having a mutation at the same position than the eight mutations (five single amino acid changes and three in-frame deletions) selected during the experimental evolution are specified. The number of exact same mutation/the total number of mutations at this specific position are specified in cells X4 to AE4.

Supplementary Table S4 (available https://research.pasteur.fr/en/project/supplementary-tables-for-biorxiv/): **Phylogroups, strain origin and enrichment in the mutations found at the 20 hotspots. A.** Phylogroup enrichment in the 361 sequences mutated at the same position than the mutants selected in our experiments as well as the 64 sequences having the exact same mutations relatively to all FimH sequences from GenBank. Ns: not significant. **B**. The 20 hotspots identified in *E. coli* FimH sequences. For each hotspot the position is specified in the third column and the first column specifies the location in the sequence. All identified alleles are listed in the fourth column with the number of sequences bearing it in the following one. The last column shows the phylogroup enrichment related to each allele when applicable. **C**. Phylogroup enrichment in the sequences bearing co-occurring mutations. The percentage of each phylogroup is specified in each case with the associated p-value between parentheses (Fisher exact test). Values corresponding to enriched phylogroups are shown in red whereas depleted ones are in blue. ns = not significant. **D.** Phylogroups enrichments in tree sub-groups. For each sub-group, an exact test of Fisher was used to identify the enriched and depleted phylogroups relatively to the reminder of the tree. The p-values were FDR corrected and a threshold of 0.05 was considered as significant.

Supplementary Table S5. (available https://research.pasteur.fr/en/project/supplementary-tables-for-biorxiv/): **Fixed-site models testing for different evolutionary pressures on the mannose lectin domain and the pilin domain. A.** Comparison between M0 and M3 models testing for selection variability among sites and M8a and M8 models testing for positive selection. lnL is the log-likelihood value for each model. ⍵0, ⍵1 and ⍵2 are the dN/dS ratios of the three site classes inferred by the M3 model and p0, p1 and p2 are the related proportions of sites in the sequence belonging to these site classes. P0, P1, P and q are the parameters estimated by the M8a and M8 models. For the BEB analysis, sites predicted to be under positive selection at the 99% level are in bold, the other are predicted at the 95% level. **B.** Branch-site model A test for episodic positive selection in sub-groups 1 to 4. lnL is the log-likelihood value for each model. In each case, the foreground branch is the sub-group under investigation and the background is the reminder of the tree. For the BEB analysis, sites predicted to be under positive selection at the 99% level are in bold, the other are predicted at the 95% level. **C.** Fixed-site models testing for different evolutionary pressures on the mannose lectin domain and the pilin domain. The first partition corresponds to the lectin domain and the second to the pilin domain. r2 is the relative substitution rate of the second partition relatively to the first partition. K1 and K2 are the transition/transversion ratio for the first and the second partition, respectively. ⍵1 and ⍵2 are the dN/dS ratio for the first and the second partition, respectively. **D.** Likelihood Ratio Test for fixed-sites models. Model A vs B tests for different substitution rates. Model B vs C tests for different substitution rates and codon frequencies. Model B vs D tests for different K and ⍵ ratios. Model B vs D2 tests for different ⍵ ratios. The resulting p-values were FDR corrected and a threshold of 0.05 was considered as significant.

Supplementary Table S6 (available https://research.pasteur.fr/en/project/supplementary-tables-for-biorxiv/): **Mutations identified in biofilm positives populations and corresponding clones from the evolution in a <u>Δ*fim*</u> genetic background at cycle 15 of the experiments**. SheetA mutations in the 3 biofilm positive populations, SheetB mutations in corresponding clones of the 3 biofilm positive populations.

For each mutation identified the gene, nature of the mutation, impact on the protein sequence as well as the frequency of the mutation in the population are shown.

**Supplementary Table S7:** *E. coli* strains used in this study.

| Strain | Genotype-Phenotype | Relevant Information | Source |
| --- | --- | --- | --- |
| MG1655 | F<sup>-</sup> lambda<sup>-</sup> *ilvG rfb-50 rph-1* | *E. coli* Genetic Stock Center GCSC#6300 | Laboratory collection |
| TG1 | F'[*traD36 proAB + lacI<sup>q</sup> lacZDM15*]<br>*supE hsdD5 thi Δ(lac-proAB)* | Strong biofilm former | Laboratory collection |
| WT *gfp* | MG1655*gfpmut3*, Km<sup>R</sup> | *gfpmut3* gene inserted at the λatt site under λP<sub>R</sub> constitutive promoter | This study |
| WT *rfp* | MG1655*mars*, Km<sup>R</sup> | *mars rfp* gene inserted at the λatt site under λP<sub>R</sub> constitutive promoter | This study |
| PcL*fimH* WT | MG1655*gfpmut3_specPcLfim*, Km<sup>R</sup>, Spec<sup>R</sup> | WT with constitutive type 1 fimbriae | This study |
| Δ*fim* *gfp* | MG1655*gfpmut3_Δfim::cat*, Km<sup>R</sup>, Cm<sup>R</sup> | WT with deleted type 1 fimbriae | Korea *et al* (1). |
| Δ*fim* *rfp* | MG1655*mars_Δfim::cat*, Km<sup>R</sup>, Cm<sup>R</sup> | WT with deleted type 1 fimbriae | Korea *et al* (1). |
| S16L | MG1655*mars_fimH_S16L*, Km<sup>R</sup> | WT with S16L mutation in *fimH*, isolated from population R4D15 | This study |
| V77A | MG1655*mars_fimH_V77A*, Km<sup>R</sup> | WT with V77A mutation in *fimH*, isolated from population R6D15 | This study |
| L79P | MG1655*mars_fimH_L79P*, Km<sup>R</sup> | WT with L79P mutation in *fimH*, isolated from population R5D15 | This study |
| G82C | MG1655*mars_fimH_G82C*, Km<sup>R</sup> | WT with G82C mutation in *fimH*, isolated from population R5D15 | This study |
| G87C | MG1655*gfpmut3_fimH_G87C*, Km<sup>R</sup> | WT with G87C mutation in *fimH*, isolated from population PS2G4 | This study |
| Δ29-40 | MG1655*gfpmut3_fimH_Δ29-40*, Km<sup>R</sup> | WT with Δ29-40 mutation in *fimH*, isolated from population G6D15 | This study |
| Δ64-70 | MG1655*gfpmut3_fimH_Δ64-70*, Km<sup>R</sup> | WT with Δ64-70 mutation in *fimH*, isolated from population G6D15 | This study |
| Δ89-139 | MG1655*mars_fimH_Δ89-139*, Km<sup>R</sup> | WT with Δ89-139 mutation in *fimH*, isolated from population R6D15 | This study |
| PcL-S16L | MG1655*mars_specPcLfim_fimH_S16L*, Km<sup>R</sup>, Spec<sup>R</sup> | S16L mutation in *fimH* and constitutive type 1 fimbriae | This study |
| PcL-V77A | MG1655*mars_specPcLfim_fimH_V77A*, Km<sup>R</sup>, Spec<sup>R</sup> | V77A mutation in *fimH* and constitutive type 1 fimbriae | This study |
| PcL-L79P | MG1655*mars_specPcLfim_fimH_L79P*, Km<sup>R</sup>, Spec<sup>R</sup> | L79P mutation in *fimH* and constitutive type 1 fimbriae | This study |
| PcL-G82C | MG1655*mars_specPcLfim_fimH_G82C*, Km<sup>R</sup>, Spec<sup>R</sup> | G82C mutation in *fimH* and constitutive type 1 fimbriae | This study |
| PcL-G87C | MG1655*gfpmut3_specPcLfim_fimH_G87C*, Km<sup>R</sup>, Spec<sup>R</sup> | G87C mutation in *fimH* and constitutive type 1 fimbriae | This study |
| PcL-Δ29-40 | MG1655*gfpmut3_specPcLfim_fimH_Δ29-40*, Km<sup>R</sup>, Spec<sup>R</sup> | Δ29-40 mutation in *fimH* and constitutive type 1 fimbriae | This study |
| PcL-Δ64-70 | MG1655*gfpmut3_specPcLfim_fimH_Δ64-70*, Km<sup>R</sup>, Spec<sup>R</sup> | Δ64-70 mutation in *fimH* and constitutive type 1 fimbriae | This study |
| PcL-Δ89-139 | MG1655*mars_specPcLfim_fimH_Δ89-139*, Km<sup>R</sup>, Spec<sup>R</sup> | Δ89-139 mutation in *fimH* and constitutive type 1 fimbriae | This study |
| *ecpR* | MG1655 *gfpmut3_Δfim::cat_ecpR_IS2(-) +5 bp*, Km<sup>R</sup>, Cm<sup>R</sup> | Δ*fim* with the IS insertion right after *ecpR* | This study |
| *ecpD/ecpR* | MG1655 *mars_Δfim::cat_ecpD_G89D_ecpR_IS2(-) +5 bp*, Km<sup>R</sup>, Cm<sup>R</sup> | Δ*fim* with G89R mutation in *ecpD* and the IS insertion right after *ecpR* | This study |

1. *Escherichia coli* K-12 possesses multiple cryptic but functional chaperone-usher fimbriae with distinct surface specificities. Environ Microbiol 12 1957-1977 doi:10.1111/j.1462-2920.2010.02202.x (2010).

**Supplementary Table S8:**
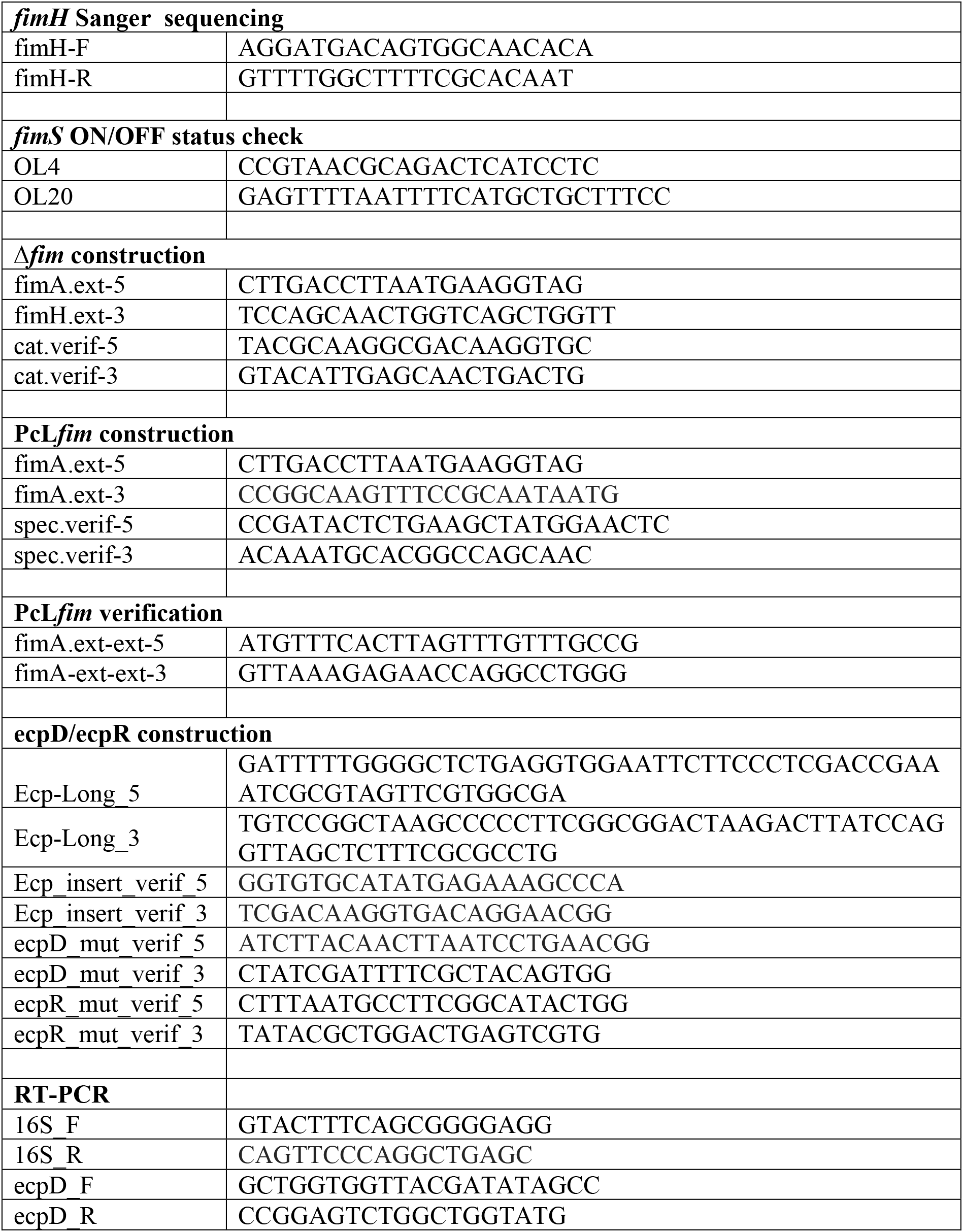
List of primers used in this study.

### SUPPLEMENTARY FIGURES

**Supplementary Figure S1.**
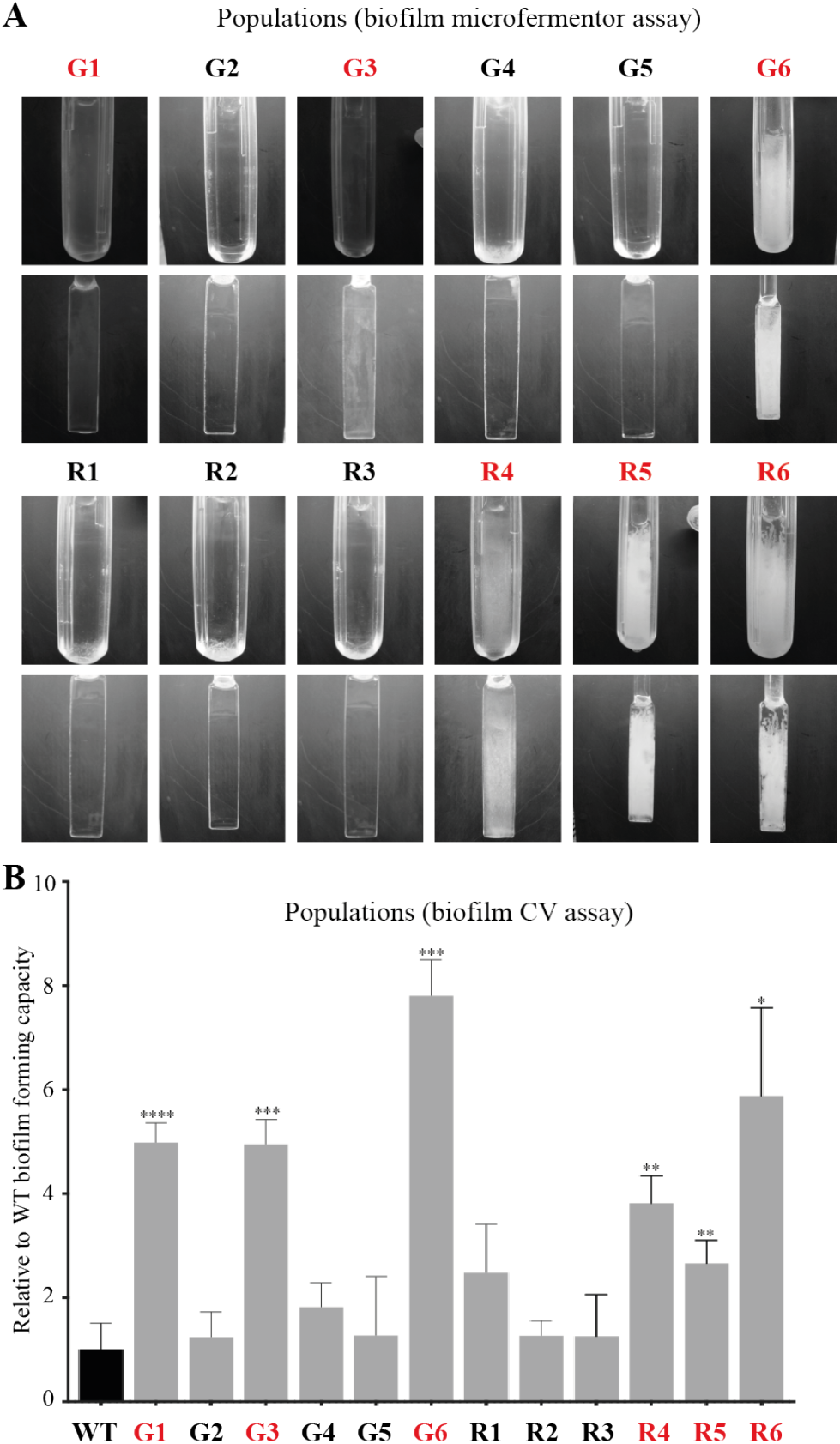
Selection of *E. coli* populations with increased biofilm capacity after positive selection. **A.** Comparison of biofilm formation on microfermenter spatulas of end-point populations after 24 hours. The bottom of the microfermenter (top) and the lower part of the spatula inserted in the microfermenter (bottom) are shown. Selected populations with increased biofilm capacity are indicated in red. **B.** Comparison of biofilm forming capacities (crystal violet CV biofilm assay) of all populations resulting from positive selection for increased adhesion at cycle 15. The ancestral wild type is shown in black. The selected populations with increased biofilm capacity are indicated in red. The relative biofilm capacities were calculated using the capacities of the parental wild type trains (set to 1). All CV biofilm assay experiments were conducted with at least 8 replicates. Statistics correspond to unpaired, non-parametric Mann-Whitney test comparing all conditions to WT. *: p<0.05; **: p<0.01; ***: p<0.001; ****: p<0.0001; absence of *: non-significant.

**Supplementary Figure S2.**
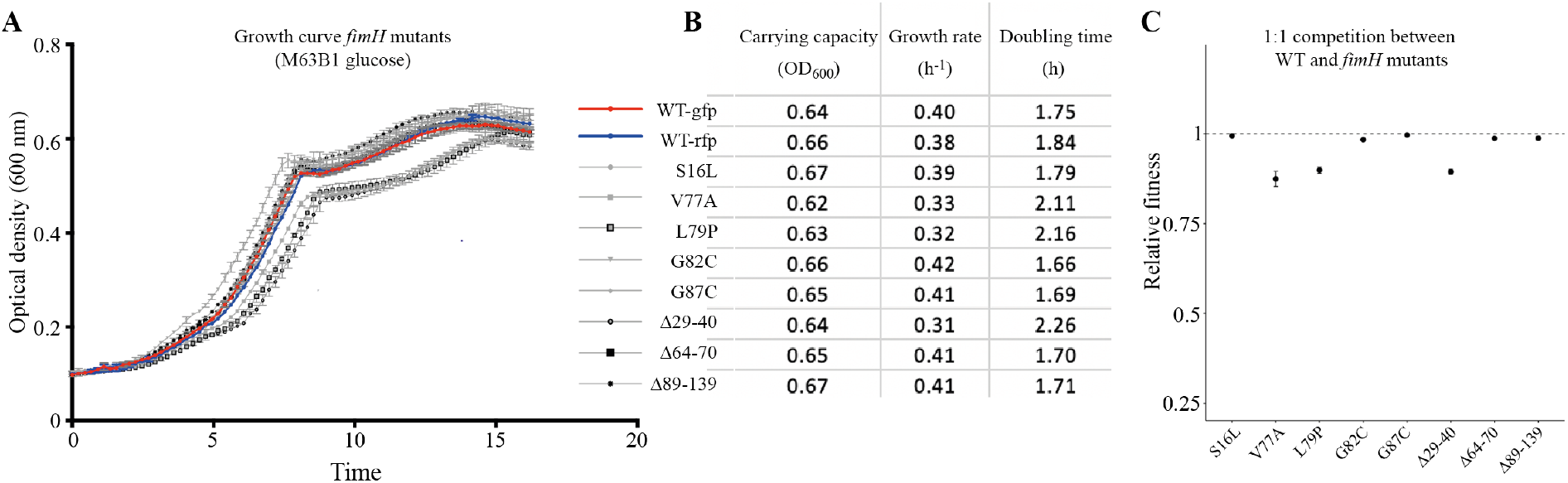
Relative growth capacities and fitness of the identified *fimH* mutants. **A**. Growth curves in minimal M63B1 glucose medium of WT ancestors (tagged with GFP or RFP) together with all identified *fimH* mutants. **B**. Carrying capacity, growth rate and doubling time of all tested strains. The growth was determined by measuring the OD600 using a plate reader. The experiment was conducted in triplicate. **C**. Competitions between individual *fimH* mutants and the WT ancestor in planktonic culture. The fitness of each mutant is expressed relatively to that of the ancestor, set to 1 (dashed line).

**Supplementary Figure S3.**
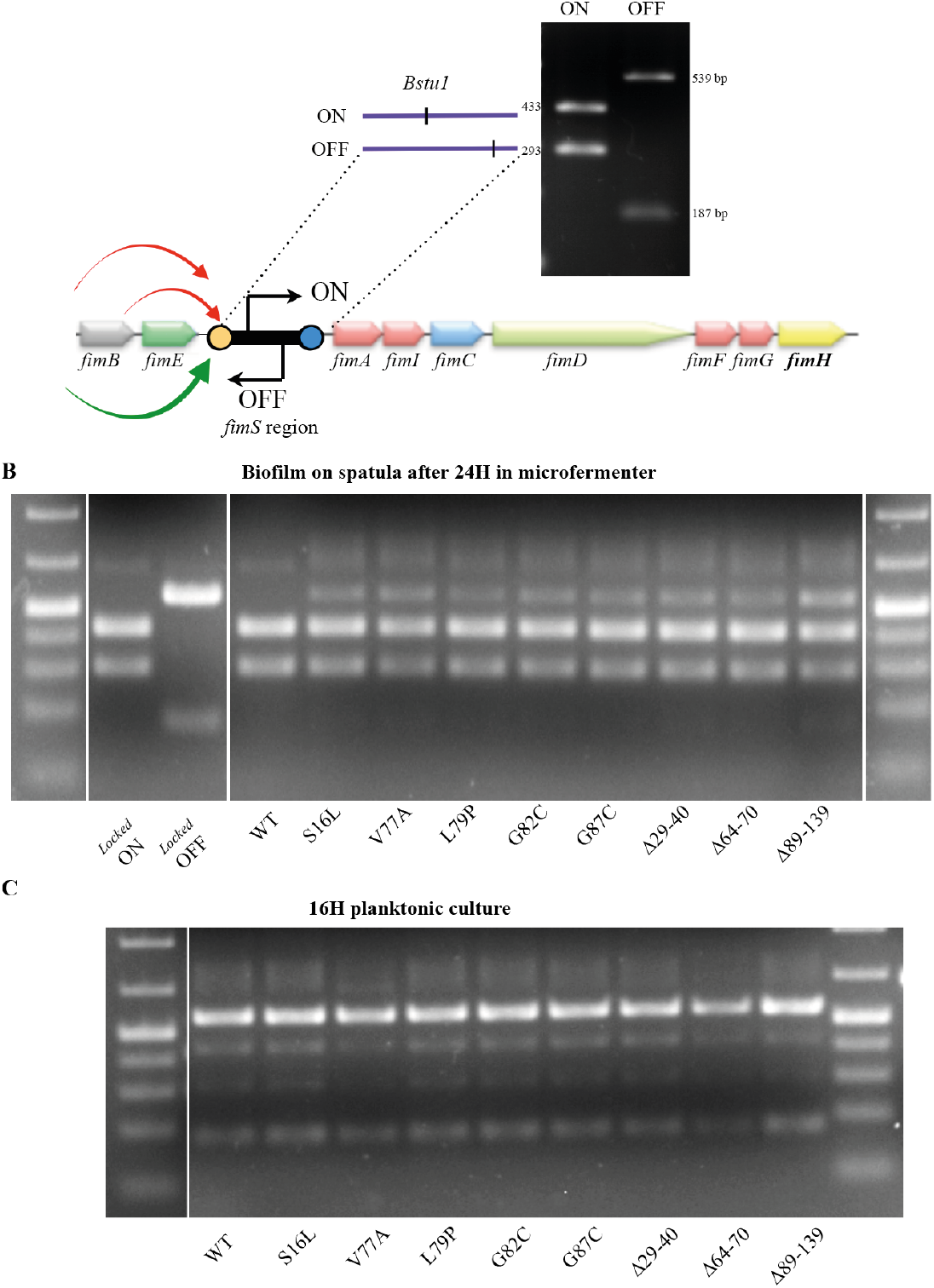
Determination of *fim* “ON/OFF” status of evolved clones. **A.** Genetic organization of the type 1 fimbriae *fim* operon and representation of the *fimS* ON-OFF orientation switch controlled by FimB and FimE recombinases. Top: PCR assay followed by BstUI restriction enzyme digestion to distinguish *fimS* ON and OFF orientation. Phase ON populations of bacteria yielded two DNA fragments 433 and 293 bp in length, whereas phase OFF populations yielded two fragments of 539 and 187 bp. **B.** PCR assay determining the *fimS* ON and OFF orientation in biofilm population on continuous flow biofilm microfermenter spatula of WT and indicated *fimH* mutants. **C.** PCR assay determining the *fimA* ON and OFF orientation in planktonic population on continuous flow biofilm microfermenter spatula of WT and indicated *fimH* mutants. Control for OFF state correspond to CJD957= VL386_*fimB*::Km locked OFF for type 1 fimbriae (1). Control for ON state correspond to CJD808= VL386_*fimB*::Km locked ON for type 1 fimbriae (2). 1. Dove SL & Dorman CJ (1994) The site-specific recombination system regulating expression of the type 1 fimbrial subunit gene of Escherichia coli is sensitive to changes in DNA supercoiling. Molecular microbiology 14(5):975-988. 2 Smith, S.G.J. & Dorman, C.J. (1999) Functional analysis of the FimE integrase of Escherichia coli K-12: isolation of mutant derivatives with altered DNA inversion preferences. Mol Microbiol 34: 965–979.

**Supplementary Figure S4.**
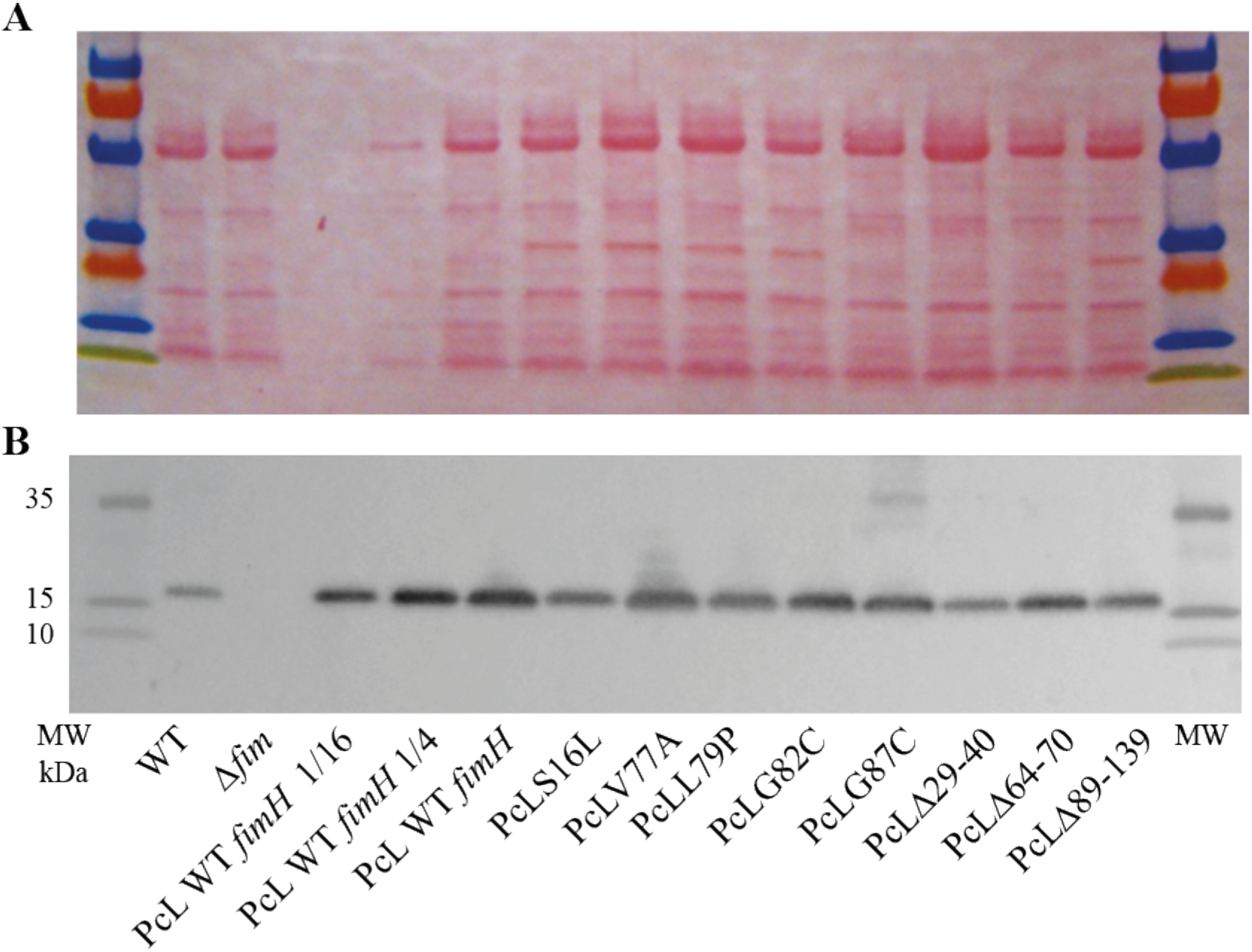
Immunodetection of type 1 fimbriae in wild type, Δ*fim*, PcL*fimH* WT, and PcL*fimH* mutant strains. **A.** Ponceau red staining of the nitrocellulose membrane after SDS-PAGE and transfer showing that OD600 = 2.0 equivalent of heat extracted proteins were loaded. **B.** Immunodetection of type 1 fimbriae (using anti-FimA antibodies) in wild type, Δ*fim*, PcL*fimH* WT, and PcL*fimH* mutant strains.

**Supplementary Figure S5.**
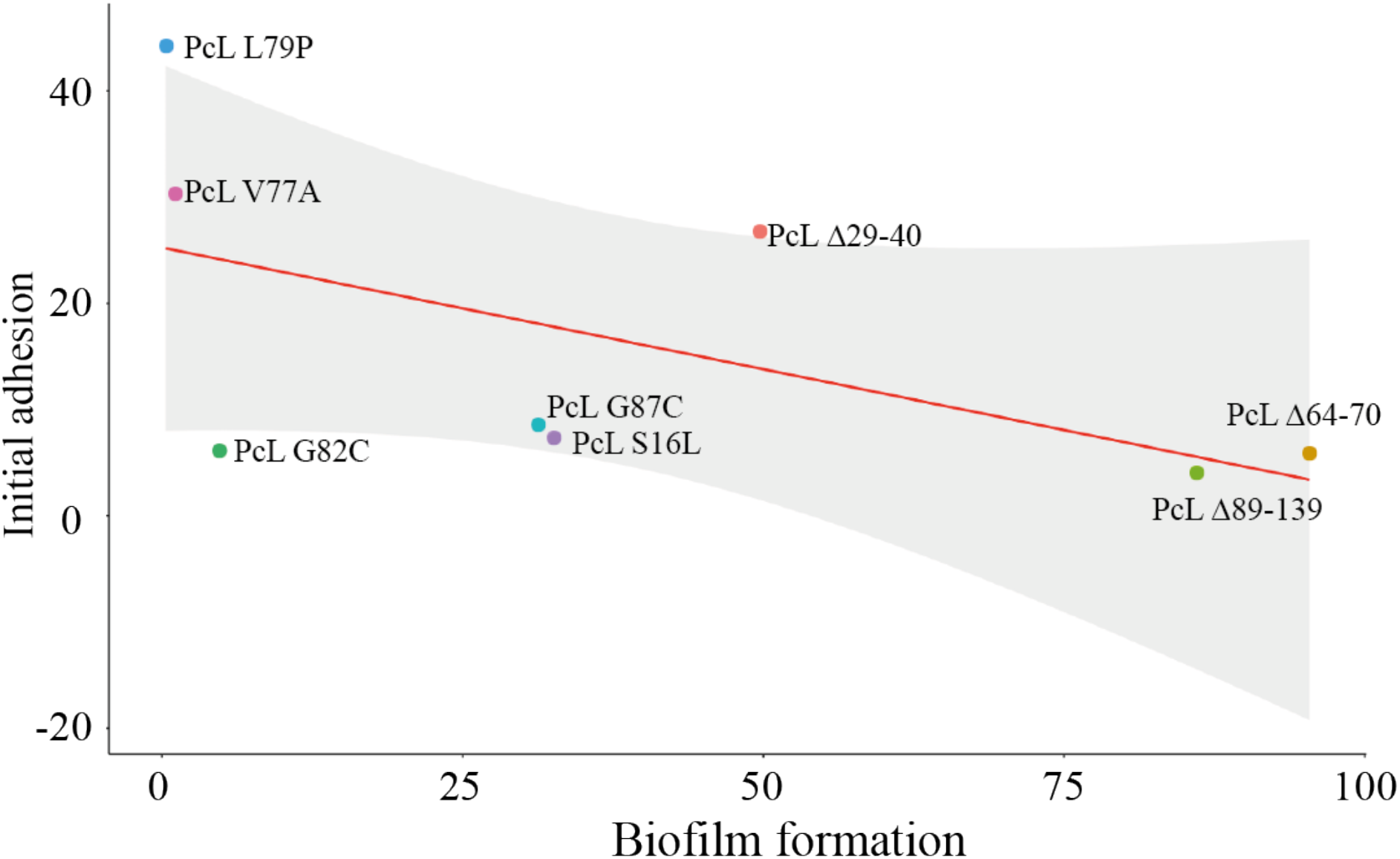
Negative trend between initial adhesion and biofilm formation in *PcLfimH* mutants. Scatter plot showing initial adhesion vs biofilm formation using data from Figure 4. The trend line is shown in red and the confidence interval in grey. Spearman’s rho = -0.76, p-value = 0.037.

**Supplementary Figure S6.**
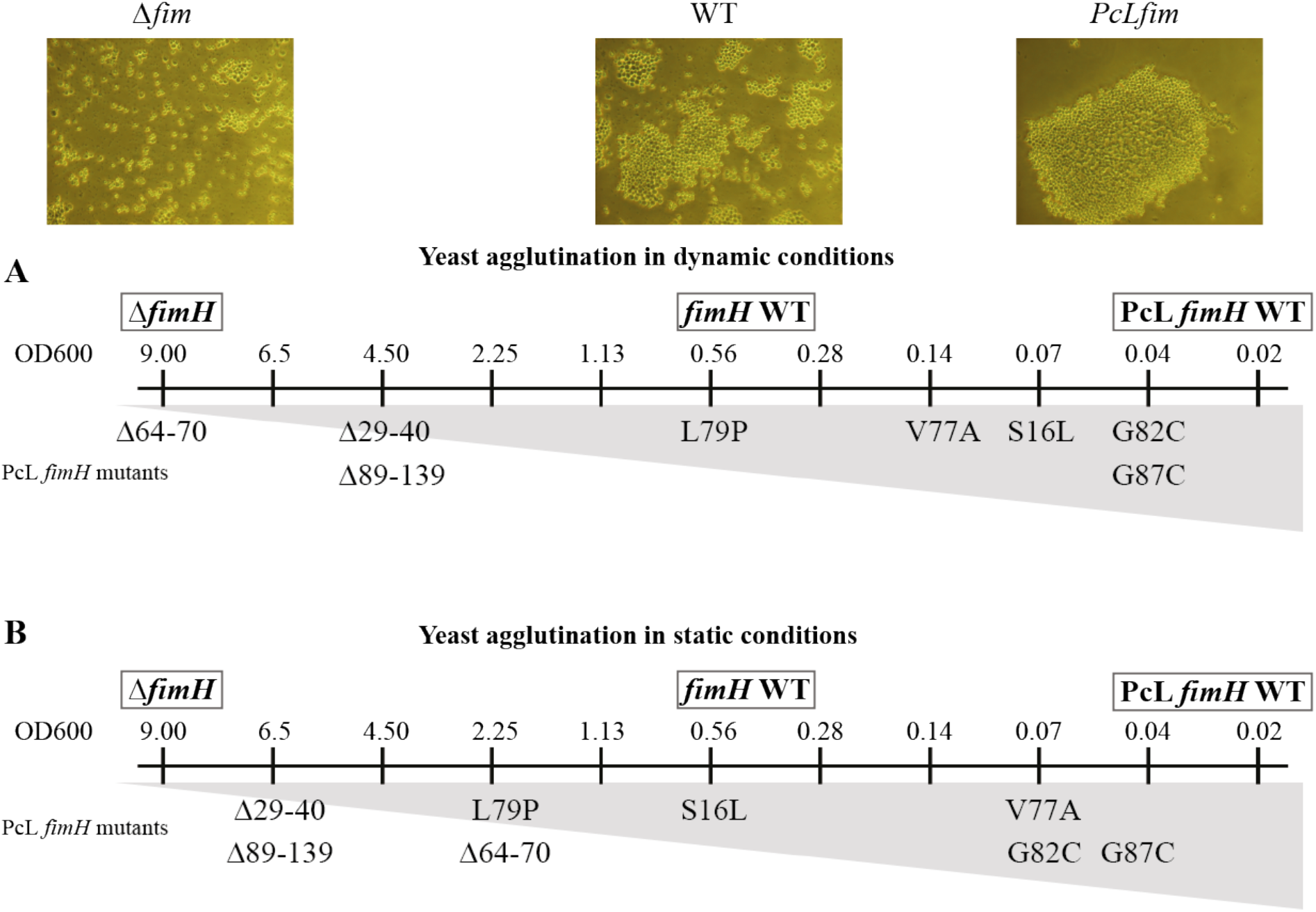
Mannose-binding capacity of *fimH* mutants. Yeast agglutination of PcL *fimH* mutants were compared with three control strains (Δ*fim*, wild type WT and PcL*fimH* WT strains) in dynamic conditions (**A)** and static conditions (**B**). The lowest bacterial concentrations (OD_600_) at which the strains agglutinate with the yeast are shown: 9.0, 0.56 and 0.04 for Δ*fim*, WT and PcL*fimH* WT strains, respectively.

**Supplementary Figure S7.**
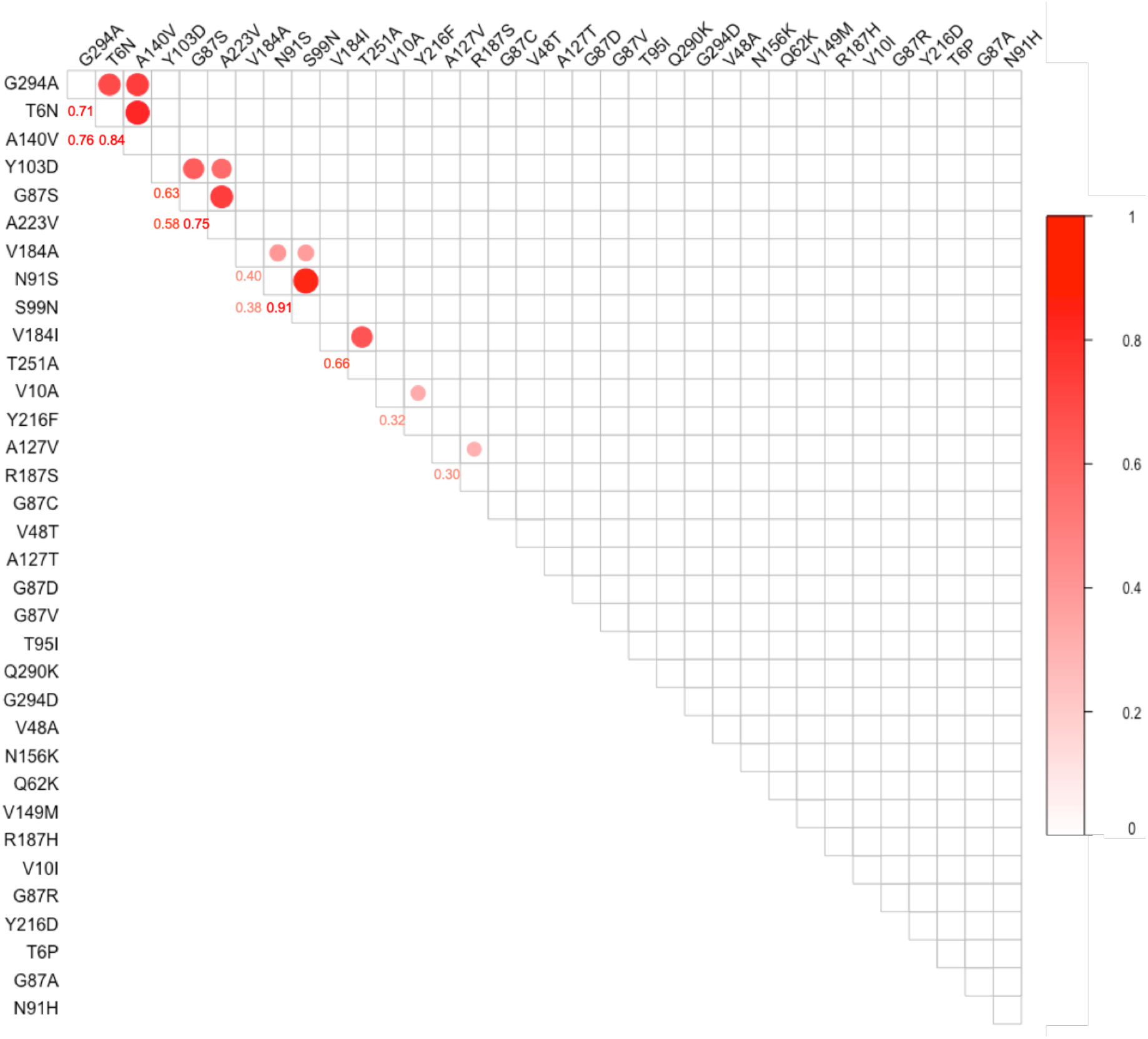
Co-occurring mutations at hotspots. All haplotypes are represented on both axes. On the upper triangle of the matrix, the correlation is represented as a circle if the correlation is statistically significant (p-value < 0.05). The color and size of the circle is proportional to the value of the correlation coefficient, which is given on the lower triangle.

**Supplementary Figure S8.**
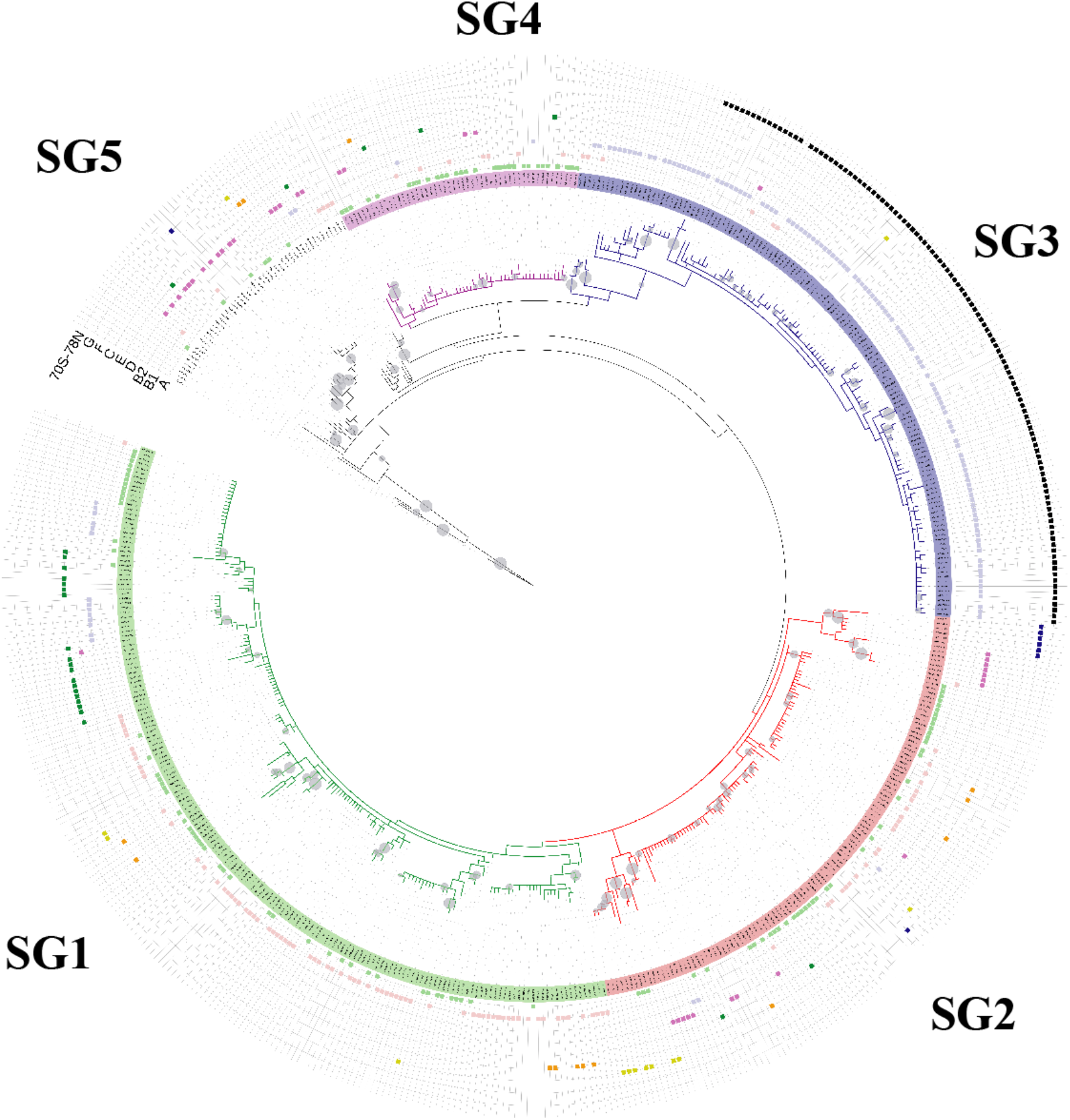
Phylogenetic tree of unique FimH sequences used for the selection pressure analysis by PAML. The tree was edited using iTOL v4 ^74^ . Bootstrap values higher than 0.5 are identified by a grey circle on the corresponding leaves of the tree. The size of the circle is proportional to the bootstrap value. The five sub-groups are represented by different colors (Sg1 in green, Sg2 in red, Sg3 in blue, Sg4 in purple and Sg5 in black). Outside the tree, nine additional information are represented. From the inside to the outside: sequences of strains belonging to phylogroup A, B1, B2, D, E, C, F and G are marked with different colored squares. Finally, sequences having 70S and 78N positions (91S and 99N in this study), which were related to UPEC strains (1), are marked with a black square. 1. Chen SL*, et al.* (2009) Positive selection identifies an in vivo role for FimH during urinary tract infection in addition to mannose binding. *Proceedings of the National Academy of Sciences of the United States of America* 106(52):22439-22444.

**Supplementary Figure S9.**
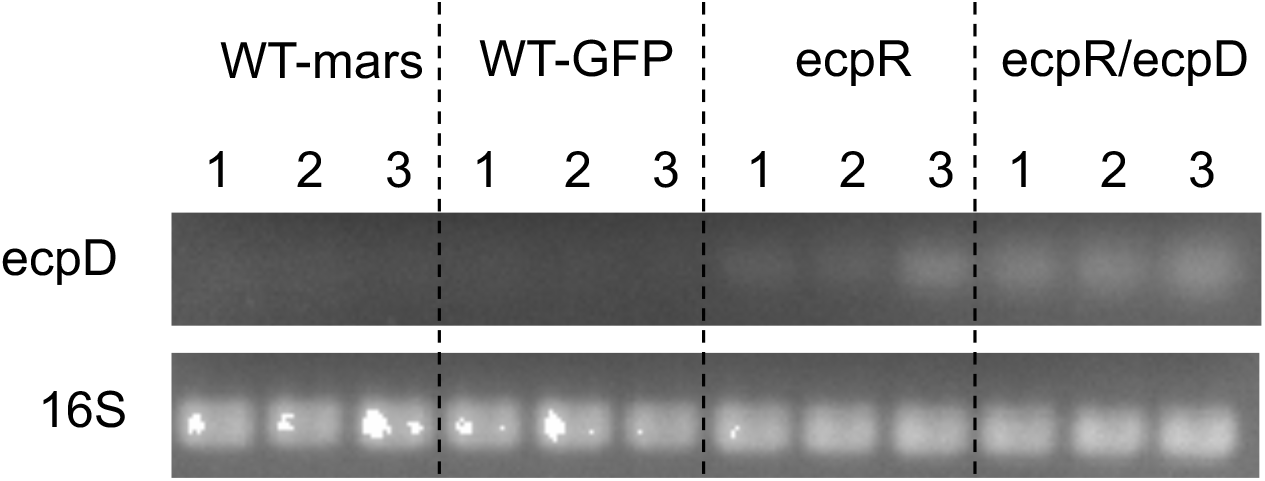
Impact of IS2 insertion on *ecp* operon expression. RT-PCR targeting *ecpD* in WT strain with either mars or GFP marker as well as in strains selected in the Δ*fim* evolution and having the IS2 insertion after *ecpR*. 16S rRNA is used as a reference. For each strain, the three biological replicates are marked with numbers.

**Supplementary Figure S10.**
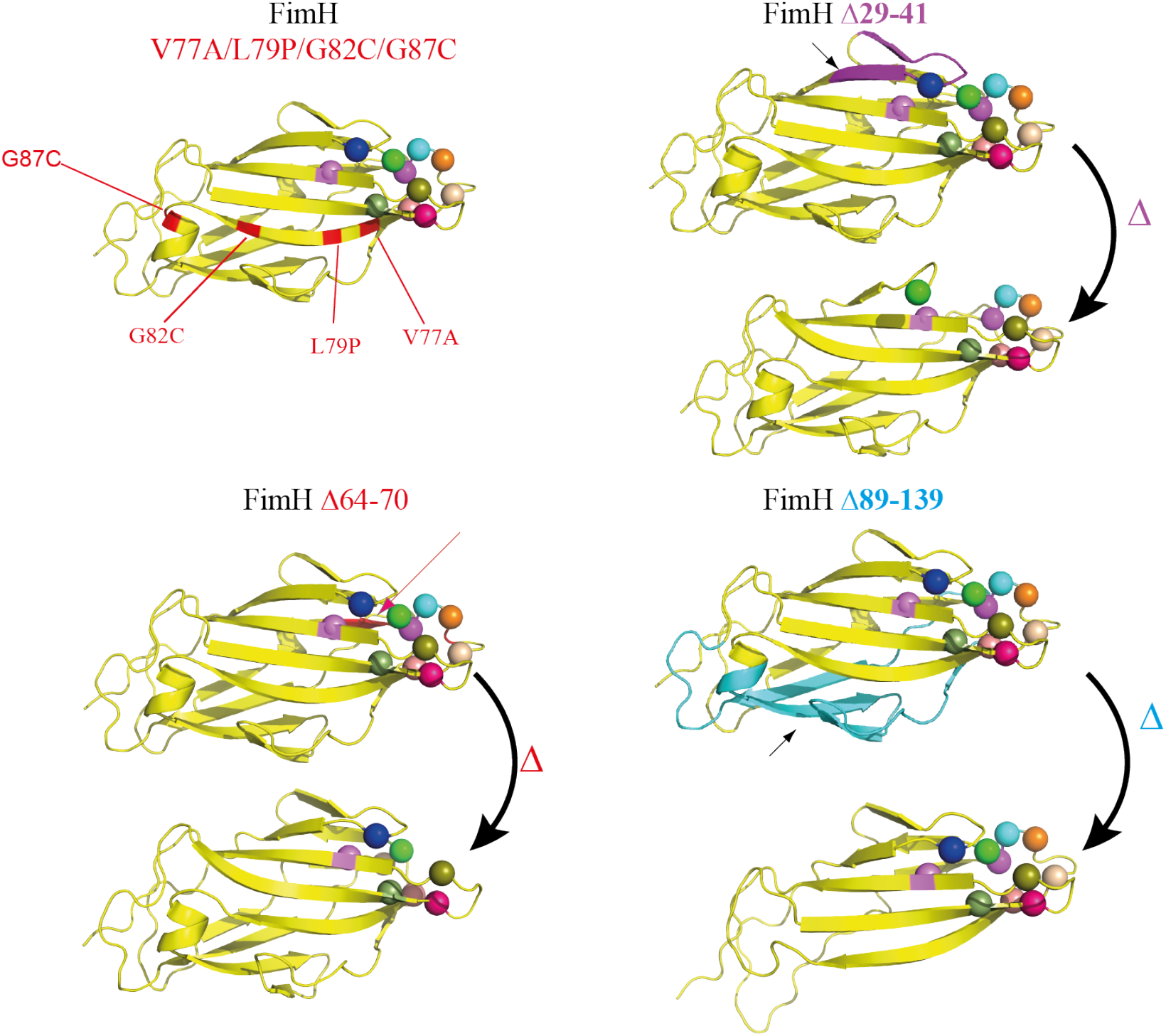
Structural changes in the FimH lectin domain with selected mutations or deletions. These positions are compared in relation to the amino-acids residues forming the mannose binding pocket and the surrounding hydrophobic ridge (represented by spheres). For the three deletions isolated in our study, the potential structural modification of the deletion have been modelled using Phyre2 (1). The modification of the presence or the organisation of these important residues is shown upon the three deletions that affected the most strongly FimH mannose-binding capacity. 1. Kelley LA, Mezulis S, Yates CM, Wass MN, & Sternberg MJE (2015) The Phyre2 web portal for protein modeling, prediction and analysis. *Nature Protocols* 10:845.

